# Dynamic refinement of behavioural restructure mediates dopamine-dependent credit assignment

**DOI:** 10.1101/2022.09.22.507905

**Authors:** Jonathan C.Y. Tang, Vitor Paixao, Filipe Carvalho, Artur Silva, Andreas Klaus, Joaquim Alves da Silva, Rui M. Costa

## Abstract

Animals exhibit a diverse behavioral repertoire when exploring new environments and can learn which actions or action sequences produce positive outcomes. Dopamine release upon encountering reward is critical for reinforcing reward-producing actions^1–3^. However, it has been challenging to understand how credit is assigned to the exact action that produced dopamine release during continuous behavior. We investigated this problem with a novel self-stimulation paradigm in which specific spontaneous movements triggered optogenetic stimulation of dopaminergic neurons. Dopamine self-stimulation rapidly and dynamically changes the structure of the entire behavioral repertoire. Initial stimulations reinforced not only the stimulation-producing target action, but also actions similar to target and actions that occurred a few seconds before stimulation. Repeated pairings led to gradual refinement of the behavioral repertoire to home in on the target. Reinforcement of action sequences revealed further temporal dependencies of refinement. Action pairs spontaneously separated by long time intervals promoted a stepwise credit assignment, with early refinement of actions most proximal to stimulation and subsequent refinement of more distal actions. Thus, a retrospective reinforcement mechanism promotes not only reinforcement, but gradual refinement of the entire behavioral repertoire to assign credit to specific actions and action sequences that lead to dopamine release.

## Background

Animals spontaneously transition amongst a repertoire of movements when exploring new environments. Movements or movement sequences that produce positive outcomes are reinforced and increase their frequency to maximize the obtaining of such outcomes^4, 5^. However, it is not completely clear how animals assign credit to the exact action that produces reward in the context of a continuous behavioral space. This credit assignment problem^2, 6–9^ during spontaneous behavior poses at least two main challenges. First, it is unclear how animals refine their behavior to preferentially perform a specific reward-producing action compared to similar actions in the behavioral repertoire. Second, it is unclear how animals derive contingency between reward-producing actions or action sequences and reward if there can be variable delays between action or sequence performance and reward delivery, with many actions interleaved.

Dopamine (DA) has been proposed to mediate credit assignment^8, 10^. At the cellular level, DA can facilitate synaptic plasticity in corticostriatal synapses^11^ within a critical time window that is behaviorally relevant^12–14^. Still, it is unknown how DA changes the dynamics of spontaneous behavior to mediate credit assignment. We therefore developed a paradigm to investigate how DA shapes the evolution of continuous behavior during action learning to gain insights into the process of credit assignment.

Conventional operant conditioning paradigms^5, 15–19^ have helped extract principles of reinforcement. However, they usually require animals to interact with devices (e.g. levers, nosepokes, joysticks) or perform a series of consummatory actions to obtain reward, all of this in specific locations in space. These aspects make it difficult to investigate how credit is assigned to a specific action or action sequence of the entire repertoire during continuous behavior. We developed a new approach to study credit assignment where we directly reinforce the execution of specific spontaneous movements by triggering DA neuron excitation and DA release upon their execution, irrespective of where in space they are executed. It combines wireless inertial sensors, unsupervised clustering of continuous behavior^20, 21^ and optogenetics^22^ in a closed-loop system linking specific action performance to immediate phasic DA release (Methods; Fig. 1a-e). This paradigm reinforces actions without requiring an animal to approach or interact with a place/object/cue, or to perform consummatory behavior.

**Fig. 1.**
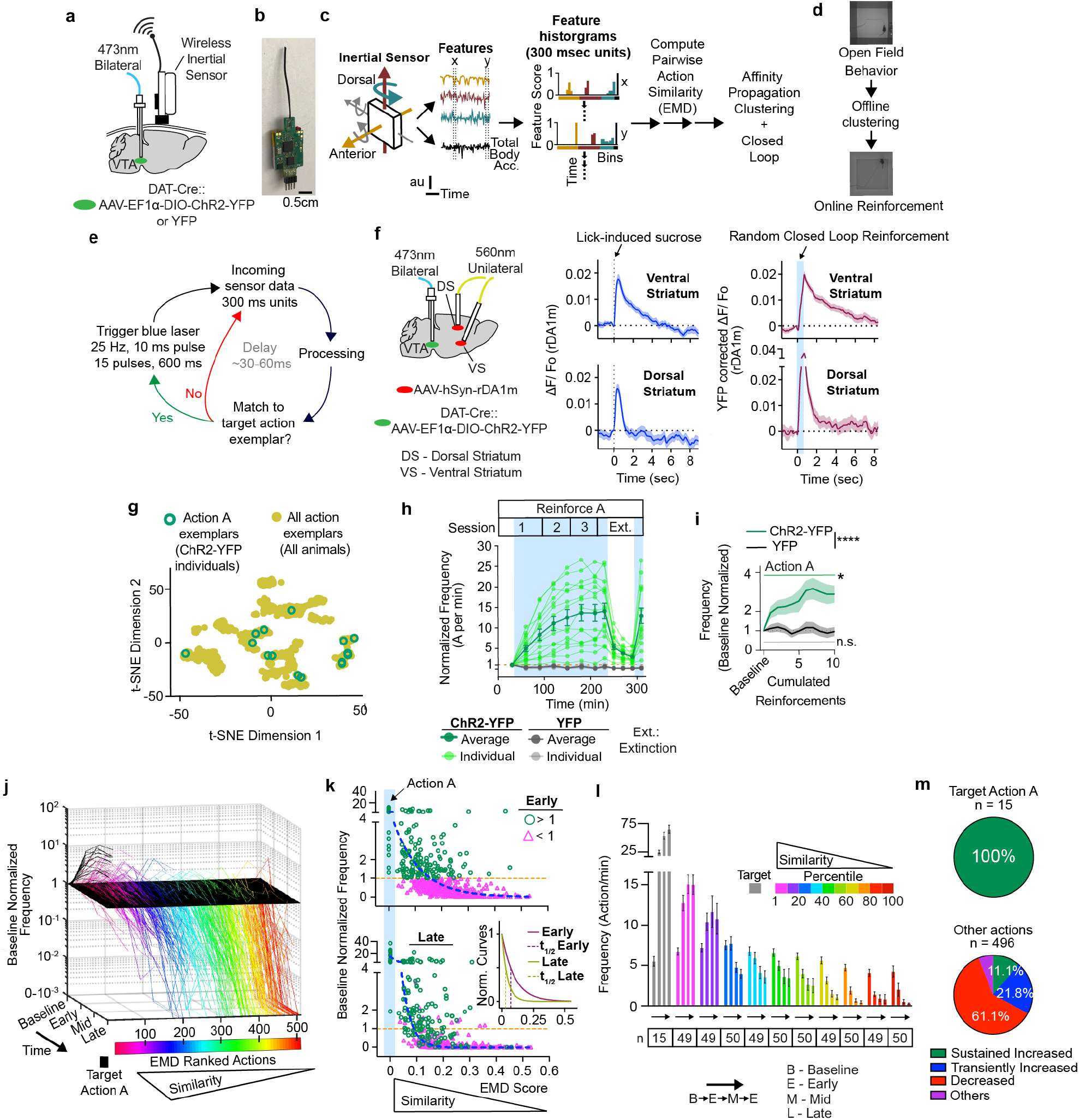
Learning of a single action from the naïve state as mediated by closed loop optogenetics. **a**, Animal implant scheme. **b**, Wireless inertial sensor. **c**, Sensor data processing. **d**, Open field behavioral clustering and action reinforcement. **e,** Closed loop schematic. **f,** Dopamine release in dorsal and ventral striatum (n = 70 sucrose rewards, 2 ChR2-YFP mice (biological replicates); n = 66 and 65 random stimulations, 2 ChR2-YFP and 2 YFP mice (biological replicates), respectively). Plots were mean, S.E.M. **g.** Action A exemplar locations in behavioral space. **h-m,** ChR2-dependent reinforcement of Action A (*n* = 15 ChR2-YFP animals (green). *n* = 10 YFP animals (grey)). Plots were mean, S.E.M. **h**, Light green/grey lines represent individual ChR2-YFP/YFP animals, respectively. **i,** Rapid increase in target action performance in response to close-loop reinforcements. Repeated measures 2-way ANOVA, significant Time x Group Interactions (F(35, 805) = 3.12; p=7.7*10^-9). Plots were mean/S.E.M. **j,** Evolution of pooled behavior repertoire (n = 509 actions, ChR2-YFP mice) across learning. **k**, Early/Late cross-sectional views of (**j**) (Early: baseline normalized frequency >1, green circles, < 1, magenta triangles). Blue dashed lines - single phase log decay fits. Bottom inset graph shows Early/Late fitted lines normalized to 1 at EMD=0. **l,** Raw frequencies across learning and target similarity percentile groups. n – sample size (actions). Plots were mean, S.E.M. Two-way mixed effects statistics in Supplementary Information. **m,** Pie chart summarizing distribution of actions according to their dynamics within reinforced Action A (left) or other actions (right). Asterisks: **** p < 0.0001. *** p < 0.001. ** p < 0.01. * p < 0.05. n.s. – not significant. See Supplementary Information for statistical/sample details.

### Rapid reinforcement of actions via closed-loop dopamine stimulation

To implement closed-loop reinforcement, we used a Cre-dependent strategy to express channelrhodopsin ChR2-YFP^22^ or the control YFP bilaterally in DA neurons of the ventral tegmental area (VTA)^25, 26^ of DAT-Cre mice^23^ (Methods; Fig. 1a, Extended Data Fig. 2a-c). We classified the entire behavioral repertoire of individual mice in a grey-walled open field (Fig. 1b-d). Self-paced behavior was monitored using a novel, wireless inertial sensor system (WEAR; Methods) that allows minimal movement restraints, high resolution behavior monitoring and fast data transmission to open-source hardware and software for online experimentation (Fig. 1b, Extended Data Fig. 1a). Affinity propagation clustering was applied to classify behavior, as it is advantageous for identifying unknown number of clusters^20, 21^, is computationally efficient^24^, and easily outputs similarity between clusters (Methods, Supplementary Methods). We identified over 30 clusters of spontaneous behavior per individual (34.3 +/- 2.1 and 35.6 +/- 2.5 actions per ChR2-YFP and YFP mice, respectively; mean +/- standard deviation,15 ChR2-YFP and 10 YFP mice), representing the entire behavior repertoire of each individual in the open field. For each animal, we chose to reinforce two clusters, or actions (hereafter named A and B), that are highly mobile and tend to have opposing feature score distributions in the A-P postural and D-V head/body turn features (Methods, Supplementary Video 1). The sampled action pairs are quite dissimilar as measured by EMD, allowing us to assess learning process for two distinct action types per animal. Variation in features across actions were allowed to extract generalizable principles common to action learning.

Although movement transitions are also accompanied by DA release in the SNc^27, 28^, and this release can pattern behavior^29^, VTA DA release upon encountering unexcepted rewards is pronounce and strongly reinforces behaviors that lead to it^29–32^. We therefore targeted VTA DA neurons and stimulated them with a pattern (25 hz, 600 ms long train) that would mimic DA release upon encountering sucrose, rather than SNc DA neurons (Supplementary Methods). Target action were different between animals and dispersed across a behavioral space (Fig. 1g; Extended Data Fig. 1-2).To evaluate whether stimulation parameters triggered DA release similar in magnitude to that triggered by sucrose reward in food restricted mice, we monitored DA release in both conditions with the GRAB rDA1m sensor^30^ in both ventral and dorsal striatum (Fig. 1f). Sucrose presentation led to a sharp increase in DA release in both areas (Fig. 1f). Interestingly, random optogenetic stimulation of DA neurons in VTA with the parameters described above, resulted in a similar phasic increase in DA not only in ventral striatum but also in dorsal striatum, although there is relatively higher DA release in dorsal striatum (Fig. 1f). This is consistent with evidence of dorsal striatum-projecting VTA neurons^28, 31^. Thus, our optogenetic stimulation triggered DA release similar in decay and spatial localization to that triggered by sucrose reward in food restricted mice (Fig. 1f), offering a suitable approach to interrogate how pairing DA release with specific action performance leads to credit assignment.

Over a 3-day, 60-90 minute/session protocol designed to probe both intra- and inter-session changes in behavior, closed-loop stimulation of VTA DA neurons upon execution of a particular target (action A) significantly increased target frequency for ChR2-YFP, but not YFP mice (Fig. 1h, Extended Data Fig.4a-b). The increased frequency of action A depends on optogenetic stimulation, as demonstrated by both extinction and re-instatement of stimulations (Fig.1h, Extended Data Fig.4c-d). During extinction, ChR2-YFP animals kept performing exploratory unrewarded bursts of action A, which could explain rapid reinstatement (Extended Data Fig. 4e,f). Just a few pairings with DA leads to rapid reinforcement, as changes in multiple parameters including decreased interval between triggers, increased action A frequency and increased average behavioral similarity towards action A become significant following 10-15 stimulations (Methods, Fig. 1i, Extended Data Fig. 5a-b).

We next examined the impact of closed loop reinforcement on non-stimulated actions. We tracked baseline-normalized frequency of all actions while sorting them based on similarity to target, using Earth-Mover’s Distance (EMD)^21, 32^ (Fig. 1j). Lower EMD value indicates increased similarity. Surprisingly, optogenetic stimulation drastically changed in the entire behavioral repertoire performed in the open field. Early on, actions most similar to target tended to also increase in frequency (Fig. 1j-l, Extended Data Fig. 5c) whereas actions most dissimilar to target tended to decrease in frequency. Repeated pairing led to refinement of actions that were performed at high frequency, and by late stages action A became the predominant action being performed, with a sharp drop-off of non-target action frequencies as similarity to target decreased (Fig. 1k-l). Such effects were not observed in YFP controls (Extended Data Fig. 5d-f). Thus, early reinforcement results in rapid reshaping of the entire behavioral repertoire, biasing animals towards actions similar to the target, and continued pairing resulted in gradual refinement and assignment of credit to the specific target action.

### Dynamics of behavioral refinement during reinforcement

To better describe individual action dynamics during reinforcement, the trajectories of all action frequency changes throughout learning were categorized (Methods; 511 actions, *n*=15 ChR2-YFP animals). Three meaningful types of trajectories were characterized by “Sustained Increase”, “Transient Increase” or “Decreased” dynamics (94% of all actions; Supplementary Methods; Fig 1m, Extended Data Fig. 6-7). The dynamics of action reinforcement were, yet again, related to the action’s similarity to target, regardless of whether actions were sorted based on their raw or percentile similarity scores (Extended Data Fig. 7b-c). Actions most similar to target were predominately Sustained Increase types. Moderately similar actions mostly comprised of Sustained Increase or Transient Increase types. Highly dissimilar actions tend to be Decreased types. Transiently increased dynamics of similar actions was not caused by stimulation of non-target actions, since mis-classification of target for stimulation was very rare (1.02*10^-5 mismatch/actual triggers, 1 in 97,924 triggers, 15 ChR2-YFP animals) and target distributions do not overlap with most other actions in the 2-dimensional t-SNE behavioral space (Extended Data Fig. 8a,b). Importantly, these transiently increased clusters show little to no overlap with target distribution^33^ and did not lead to DA triggers (Supplementary Notes, Extended Data Fig. 8c). Thus, target misclassification was not a major cause of transiently increased action dynamics. These results show that the dynamics of action reinforcement are greatly related to the similarity to target action, in that animals initially increase reentrance of similar actions before homing in on target.

### Reinforcement and refinement after reversal of action-reward contingencies

Next, we asked if animals could follow changes in contingency between action and closed-loop DA stimulation. We therefore chose a different action, action B, which is clearly distinct from the action A for each animal (Methods, Fig. 2a, Extended Data Fig. 1c) and started delivering DA stimulation after action B. Chosen action A/B pairs were relatively dissimilar in the context of entire action similarity distributions (Fig. 2b). Upon reinforcement, previously trained ChR2-YFP, but not YFP animals increased action B performance over time and reduced action A frequency to baseline levels (Fig. 2c-e, Extended Data Fig. 9). Maintenance of action B performance depended on continual reinforcement (Extended Data Fig. 9f-g). Similar to action A, action B credit assignment unfolds by initially biasing the entire repertoire towards performance of actions similar to target B and away from dissimilar actions. This was again followed by gradual refinement for action B relative to similar actions over pairings (Fig. 2d-e, Extended Data Fig. 9h). We confirmed that action learning is contingent on action B happening before reinforcement by degrading the contingency between action B and DA, by giving random stimulations in the same average frequency but unpaired to Action B; the contingency was re-instated upon resuming action B-stimulation pairings (Fig. 2f, Extended Data Fig. 9i). Although similar patterns of behavioral refinement were observed for actions A and B, differences in the initial responses were noted (Fig. 2g-j; Supplementary Notes). Still, animals can follow changes in the contingency between actions and DA release and assign credit to a new action through a similar process of behavioral repertoire refinement.

**Figure 2.**
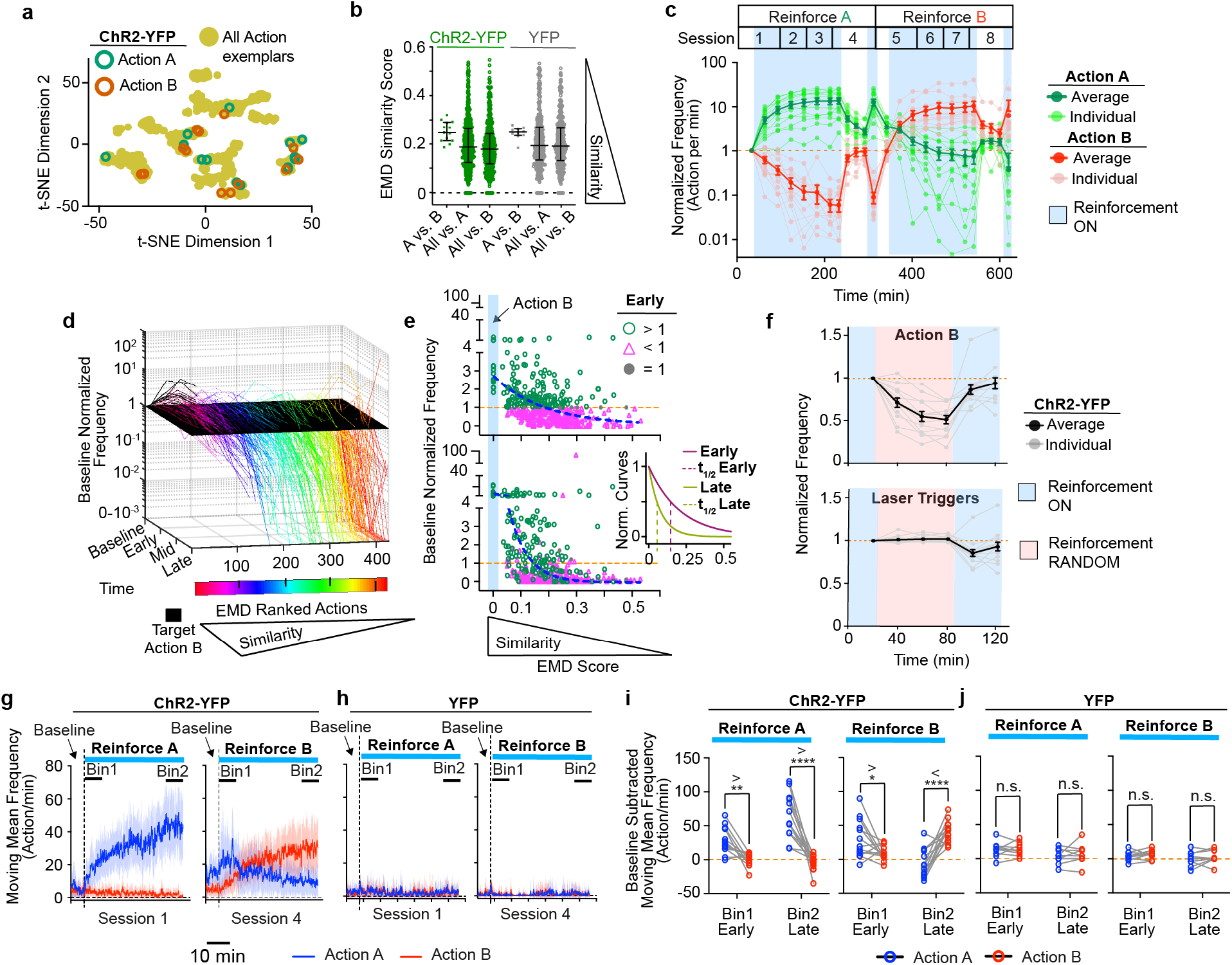
Transitioning from learned action to reinforcing new action. **a-j,** Animals (biological replicates) reinforcing for Action A (*n* = 15 ChR2-YFP) to Action B (*n* = 13 of 15 ChR2-YFP). *n* = 10 YFP All *n* indicates animals - biological replicates. **a.** Action A and B exemplar locations in behavioral space. **b,** Action similarity comparisons (A vs. B; *n* = 13/10, ChR2-YFP/YFP; All vs. A; *n* = 514/356, ChR2-YFP/YFP) or Action B (All vs. B; *n* = 443/356, ChR2YFP/YFP). Plot indicates median/interquartile range. **c,** Reinforcement for Action A and B in ChR2-YFP animals. Plot indicates mean/S.E.M. **d,** Evolution of pooled action repertoire (*n* = 427 ChR2-YFP actions) reinforced for Action B. **e,** Early/Late cross-sectional views of (**d**). Blue dashed lines indicate fitted decay curve. Bottom inset graph shows normalized Early/Late fitted curves. **f,** Contingency degradation of Action B. Target random laser triggers frequencies (bottom) is based on initial Action B performance prior to contingency degradation. Plots indicate mean/S.E.M. **g-j,** Action A (blue) induced by reinforcement for Action B in experienced ChR2-YFP animals. **g-h,** Moving mean frequencies over reinforcement for Action A or B. Dashed, vertical line mark first reinforcement. Plots are mean/S.E.M (colored fill). Bin1/Bin2 are time bins for (**i-j**). **i-j,** Frequency measures within time bins noted in **(g,h).** Repeated measures two-way ANOVA reveal significant difference across time and actions A/B frequencies (ChR2-YFP: Reinforce A, F(1,24)=34.4, p=4.8*10^-6; Reinforce B, p=1.2*10^-8). Two-sided Šidák’s post hoc multiple comparisons test. Asterisks except in (**h**): **** p < 0.0001. ** p < 0.01. * p < 0.05. n.s. – not significant. See Supplementary Information for statistical/sample details.

### Temporal constraints of DA-dependent reinforcement

Reinforcement is thought to occur on behavior that precedes reward in time^10, 12, 13, 19^, and temporal contiguity between action and reinforcement has long been recognized^34–36^ and observed above (Fig. 2f). Therefore, we investigated if in addition to behavioral similarity, the temporal relationship between action and stimulation influenced the dynamics of behavioral repertoire evolution during reinforcement and credit assignment.

The median inter-target action interval decreased with stimulation in ChR2-YFP mice (Methods; Fig. 3a,b). We therefore examined the distribution of the action dynamic types categorized above (Extended Data Fig. 7a) according to both 1.) an action’s similarity to target and 2.) the median time of that action’s performance leading into target during baseline, before reinforcement (Fig. 3c-e). These two dependent variables were not significantly collinear (Methods). Multinomial logistic regression showed that action similarity and baseline temporal proximity to target together predict action dynamic type upon reinforcement better than either factor alone. (Fig. 3f,g; Methods; Supplementary Methods and Table). Thus, DA reshapes behavioral repertoire by reinforcing both actions similar to the target action and actions that happen to be performed temporally close to the reinforcer, as suggested before^13^.

**Figure 3.**
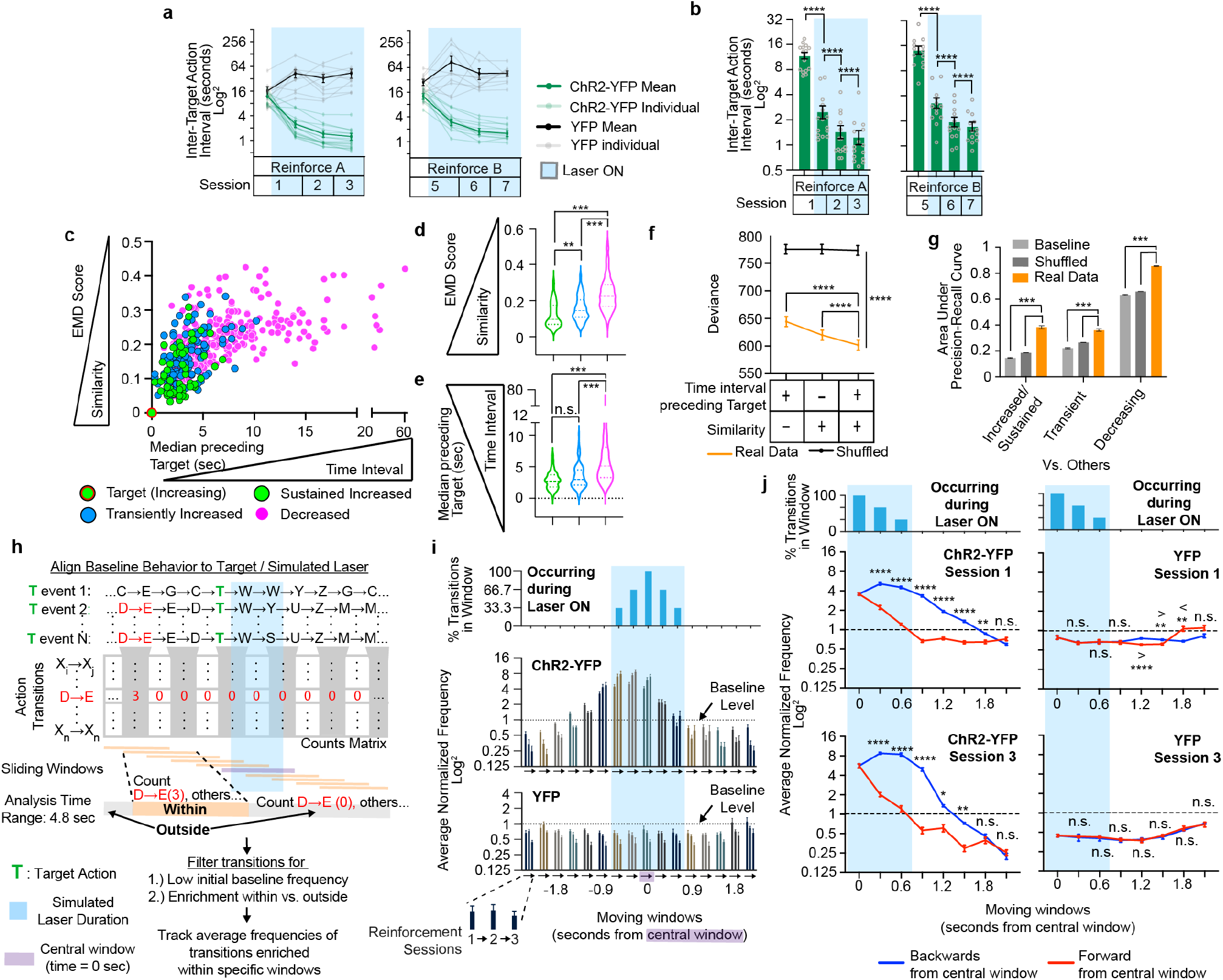
Dopamine mediates retrospective reinforcement of freely moving behavior. **a-b**, ChR2-dependent reinforcement decrease inter-action intervals for Action A (*n* = 15 ChR2-YFP) and B (*n* = 13 of 15 ChR2-YFP). *n* = 10 YFP. (n: animals - biological replicates). Plots are mean/S.E.M (**a-b**). Significant difference across time and ChR2-YFP/YFP (Mixed Effect Model. Action A: F(3,69) = 72.26, p=3.0*10^-21. Action B: F(3,62) = 33.78, p=4.6*10^-13) **b**, Post-hoc 2-tailed Tukey’s multiple comparisons of (**a**). **c-d**, Distribution of action dynamic types (*n* = 464 non-target actions, 15 target actions, 15 ChR2-YFP animals) according to target similarity (**c,d**), median time to target (**c,e**). **d-e**, Violin plots show median/quartiles. Two-tailed permutation tests with Bonferroni-adjusted p-values. **f-g**, Multinomial logistic regression of all factor combinations in Real data (200 independent models) versus Shuffled data (10,000 independent models, 50 independently shuffled datasets). Baseline – 200 independent models. **f.** 2-factor model fits data better than 1-factor models. Groups differ across combinations (repeated measures, two-way ANOVA. F(2,30594) = 1082, p=0.0*10^0). 2-tailed post-hoc Dunnett multiple comparisons test. Plots are mean/s.t.d. **g**, Performance of double-factor regression model measured with area under the precision-recall curves (AUPRC). Two-tailed permutation test with Bonferroni-adjusted p-values (p=5.9*10^-4 all comparisons). Plots are mean/S.E.M. **h**, Pipeline for identifying sliding window-enriched action transitions. **i.** ChR2-dependent reinforcement for Action A increases sliding window-enriched action transitions prior to and during stimulation. The average normalized frequency of action transitions enriched within specific sliding windows are plotted over Sessions 1-3. Top plot, percent of transitions occurring during stimulation in each sliding window. Middle and bottom plots, mean/S.E.M. **j**, Quantification of (**i**), mean/S.E.M. Significant difference across time and Retrospective/Forward reinforcement directions (Mixed Effect Modeling. ChR2-YFP Session1: F(6,168) = 114.8, p=8.7*10^-57. ChR2-YFP Session 3: F(6,168) = 46.62, p=2.5*10^-33, YFP Session1: F(6,108) = 10.52, p=3.6*10^-9. YFP Session 3: F(6,168) = 0.8992, p = 0.49). Post-hoc two-sided Šidák multiple comparisons. **** p < 0.0001, *** p < 0.001, ** p < 0.01, * p < 0.05, n.s. – not significant. See Supplementary Information for statistical/sample details.

To more rigorously test whether DA reinforcement acts in a retrospective or prospective manner, we refined our analysis by examining 1^st^ order action transitions leading into and out of stimulation (Methods; Fig. 3h-j). Action transitions enriched within specific 1.2 second sliding windows were analyzed to distinguish more clearly behavior that occurred leading up to, during, and after DA stimulation (Methods). We identified baseline-occurring action transitions enriched within specific sliding windows centered around target action and tracked their average frequencies per window over the course of closed loop reinforcement. Action transitions enriched in windows up to 1.2 seconds prior to stimulation onset, as well as during stimulation, were reinforced early on (Fig. 3i). However, action transitions enriched in windows following stimulation were not reinforced, suggesting an asymmetric process. Indeed, action transitions enriched in windows leading into stimulation were also preferentially reinforced over those enriched in windows after stimulation (Fig. 3j). Thus, DA stimulation promotes reinforcement of behaviors occurring during stimulation and a few seconds before stimulation.

### Credit assignment for action sequences

In the real world, when animals are spontaneously shifting between actions in their repertoire, outcomes are often not the result of a single action but rather of a sequence of actions performed at variable time intervals, and with other actions interleaved. We therefore investigated the dynamics of reinforcement when release of DA was contingent upon performance of a sequence of two target actions, target actions 1 (T1) and 2 (T2), where variations in time interval between the two targets, as well as interleaving actions were allowed. We applied closed loop optogenetics to ask whether naïve animals can learn a T1➔T2 reinforcement rule, where the delays between T1 and T2 are governed by the spontaneous behavior of the animals and not experimentally controlled (n=15 ChR2-YFP and 10 YFP mice, Fig. 4a, Extended Data Fig. 2b, 3a,d-e, 10-14). Various T1/T2 pairs were sampled, with focus on sequences sharing general commonalities in movement order across animals (Methods; Extended Data Fig. 1d,f-g). Overall, mice learned to increasingly perform these sequences to obtain DA stimulation. Some animals showed a ChR2-dependent increase in reinforcement within 5 sessions, but others experienced a lag in learning (Fig. 4b). We hypothesized that this relates to the initial time distance between T2 trigger and the closest distal T1 (T1➔T2 interval). Indeed, animals reinforced for action pairs with initially long interval values tended to learn slower (Fig. 4c-d). To capture a time point whereby individuals reach similar rising phase in their respective learning curves, a criterion frequency was set (Methods). 14 of 15 trained animals eventually reached criterion (Fig. 4e; Extended Data Fig. 10). Sequence performance depended on continuing DA pairings (Fig. 4f,g). Learning was also revealed by decreases in the median T1➔ T2 time intervals (Fig.4h-i) and convergence of T1-to-T2 frequency ratio towards 1 (Fig. 4j). To quantify the specific credit assignment of T1 and T2 we used a refinement index that compares the median frequency of actions uniquely similar to T1 with those uniquely similar to T2, with the frequencies normalized by either that of T1 or T2 (Methods). This index is based on the observation that actions most similar to target decrease in relative performance over time (Fig. 1k, inset). Values below 1 indicate greater refinement. By the end of learning, T1 and T2 became credited as the reward-producing actions relative to their similar counterparts (Fig. 4k). YFP controls did not show learning trends (Fig.4d-e,4h-i). Thus, closed loop reinforcement promoted learning of a two-action sequence rule in freely moving mice starting from a naïve state.

**Figure 4.**
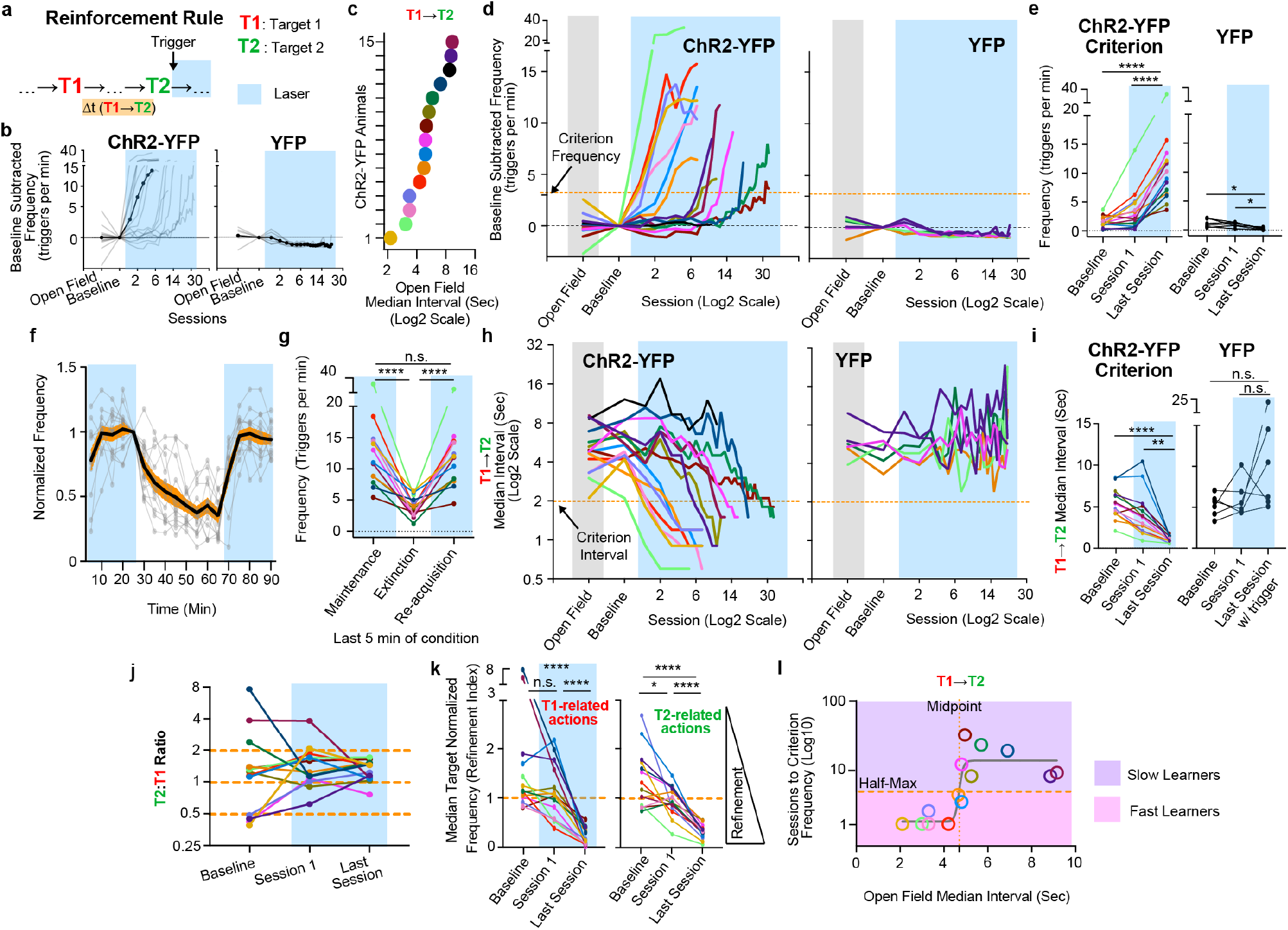
Relationship between pre-reinforcement inter-action intervals and learning of a two-action sequence. **a**, Schema. **b-l,** *n* = 15 (**b,d,h**), 14 (**e,i-l**) or 13 (f,g) ChR2-YFP, 6 YFP animals (biological replicates). Repeated measures one-way ANOVA, post hoc Šidák tests applied in (**e**,**g**,**i**,**k**). Plots of individuals in (**d-e**). **b**, ChR2-dependent increase in T1➔T2 triggers (no laser during open field / baseline). **c**, Open field inter-action intervals of T1/T2 pairs chosen. Same color codes in (**d,h**). **d**, Individual learning curves labeled by color codes in (**c**). **e**, Frequency changes over conditions. Repeated measures One-way ANOVA (F(1.911,24.85)=51.02, p=2.2*10^-7). **f-g**, Extinction of T1➔T2 sequence (ChR2-YFP). **f**, Plot shows mean(black)/S.E.M.(orange fill)/individuals(grey). **g,** Frequency changes over extinction conditions Repeated measures One-way ANOVA (F(1.073, 12.87) = 52.96, p=9.8*10^-6). **h-i**, ChR2-dependent decrease in T1➔T2 intervals. Repeated measures One-way ANOVA (F(1.377, 17.90) = 35.95, p=1.5*10^-5) (**i**). Log2 y-axis was used to help visualize interval changes in animals starting with lower initial values **j**, T2:T1 frequency ratios (ChR2-YFP) **k**, Target refinement shown by median target normalized frequencies of related actions. Repeated measures One-way ANOVA (T1: F(1.237, 16.08) = 43.38. T2: F(1.171, 15.22) = 48.74. Both p=4.4*10^-6). Individual color code as in (c,g). **l**, Sigmoidal relationship between open field T1➔T2 interval and sessions to criterion frequency. Log10 y-axis was used to capture the relationship between large values and smaller values in the same visualization.

Importantly, the initial median T1➔T2 interval of action pairs was inversely related to the eventual number of sessions required for each ChR2-YFP animal to reach criterion frequency (Fig. 4l). A sigmoidal curve was fit to the data, indicating that initial intervals longer than the sigmoidal midpoint associated with slower learning (Fig. 4l). ChR2-YFP animals were divided according to the half-maximum point of the sigmoidal curve into ‘Fast Learners’ and ‘Slow Learners’. Fast Learners quickly reached criterion frequency and low T1➔T2 time intervals, whereas Slow Learners were delayed in reaching criterion frequency and low T1➔T2 intervals. Slow Learners tended to suddenly increase sequence frequency in sessions that showed a drop in the median T1➔T2 interval to below 2-4 seconds (Fig.4d,h). In contrast, there was no stable sigmoidal relationship between T1-T2 action similarities and sessions to criterion frequency (Extended Data Fig. 11b). Further, no relationship existed between baseline frequency or initial inter-trigger intervals and sessions to criterion (Extended Data Fig.11c-d). Importantly, the observed patterns held when we analyzed learning by matching number of reinforcements (Extended Data Fig. 11e-g), indicating that they were not caused by Fast learners having more stimulations/reinforcers. Lastly, we evaluated whether differential conditionability^37–40^ account for sequence learning differences(Extended Data Fig. 12). Target actions showing less conditionability in single action reinforcement did not differ in initial baseline frequencies but tended to have more action types transitioning into and out of them at baseline (Extended Data Fig. 12d-f). Thus, differential conditionality amongst target actions relates to greater variation in behavioral environment surrounding target. However, the same parameters do not account for variation in learning rate across animals in action sequence reinforcement experiment (Extended Data Fig. 12g-l). These results support the idea that the initial median time distances between distal action T1 and proximal action T2(which produced DA stimulation) modulated how fast animals learned to effectively perform the reinforced action sequence.

If DA retrospectively reinforces actions performed earlier in time, the action most proximal to reinforcement, T2, should experience earlier refinement relative to the distal action, T1. We again used the median target normalized frequencies of actions uniquely related to T1 or T2 as refinement indices (Methods). T2 clearly refines towards its most refined level earlier than T1, at least in some animals (Fig. 5a). We calculated differential refinement between the two actions by subtracting the area under the T1 refinement curve from that of T2. Positive values indicate differential refinement favoring T2, and vice versa. Open field median T1➔T2 interval was linearly related to differential refinement between T1 and T2 (Fig. 5b). This trend holds even when accounting for within-session refinement (Methods, Extended Data Fig. 13a). Thus, for longer T1➔T2 median intervals, T2 spends more sessions being relatively more refined than T1, and this pattern cannot be explained by other potential covariates: 1.) initial intervals between proximal action and the next initiation of sequence (T2➔T1), 2.) similarity between T1 and T2 (Fig. 5b, right graph, Extended Data Fig. 13b).

**Figure 5.**
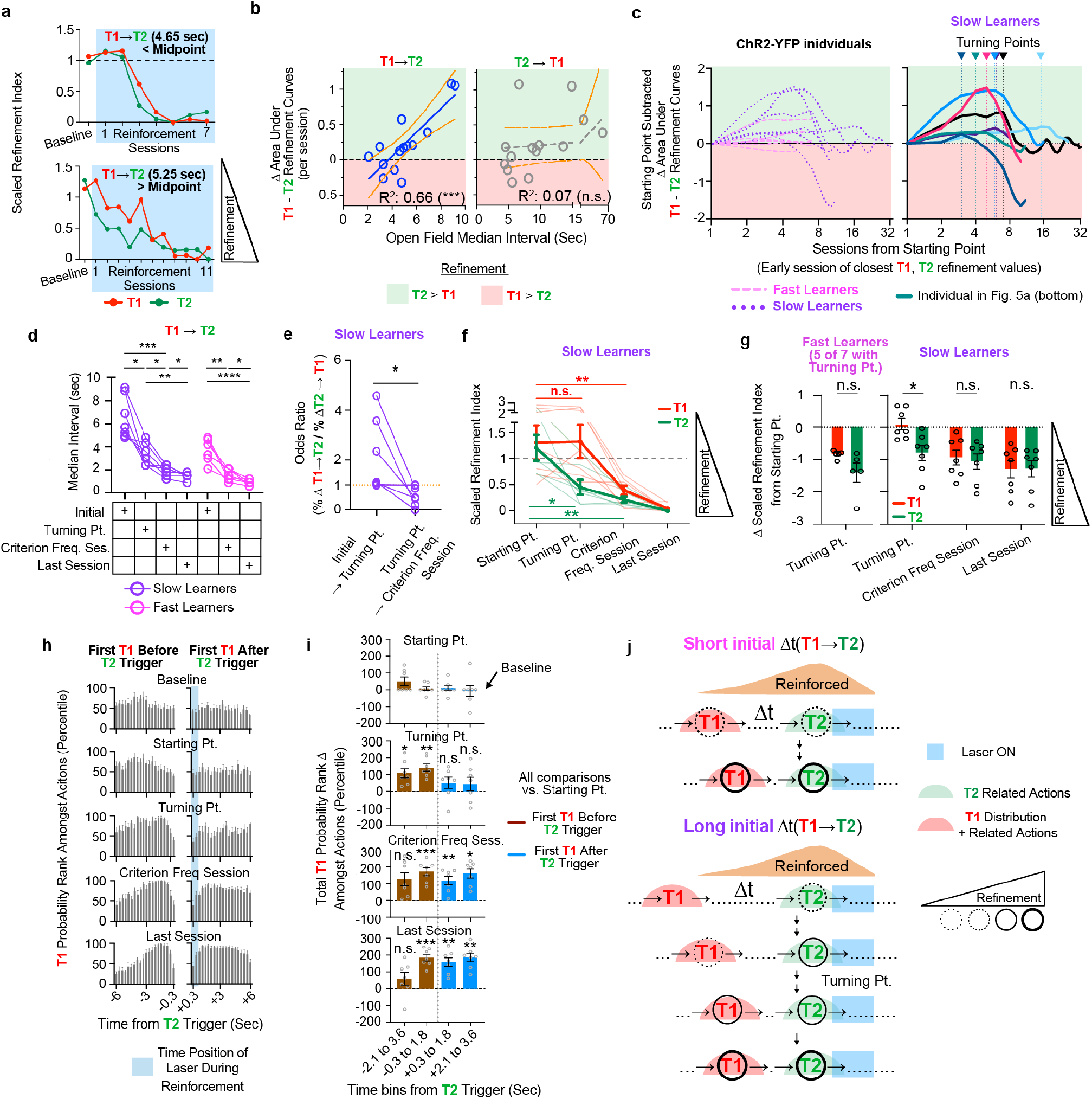
Behavioral process underlying learning of a two-action sequence. *n* =14 ChR2-YFP (7 Slow, 7 Fast Learners) (*n*: animals - biological replicates)**. a**, T1/T2 refinements in two ChR2-YFP individuals. **b**, Linear relationship between initial T1➔T2 interval and differential T1-T2 refinement. F-test: Non-zero slope significance: T1➔T2, p = 0.0004, T2➔T1, p = 0.7063. **c**, Progression of differential T1-T2 refinement from Starting Point in Individual Slow Learners. **d,** T1➔T2 interval significantly decreased by Turning Point in Slow Learners. Repeated-measures 2-way ANOVA: time-specific difference (Slow Learners – F(2.184, 26.20)=54.21, p=5.3*10^-10;Fast Learners – F(1.700, 20.40)=92.12, p=6.3*10^-9). Post hoc 2-tailed Tukey’s multiple comparisons test. **e,** Odds ratio of T1➔T2 / T2➔T1 interval changes. 2-tailed, paired Wilcoxon test (p = 0.0312, *n* = 7 animals). **f,** Preferential refinement of T2 relative to T1 by Turning Point in Slow Learners. Raw scaled refinement indices. Repeated measures, mixed effects model. Significant main effects. Time (F (2.184, 26.20) = 54.21, p < 0.0001). Post-hoc 2-sided Šidák multiple comparisons test. **g,** Starting Point-subtracted scaled refinement indices. Left: Fast learners. Welch’s 2-tailed t-test. Right: Repeated measures, mixed effects model. Significant difference between Time and Group (F(3,36) = 4.276, p = 0.011). Post-hoc 2-tailed, Šidák multiple comparisons test. **h,** Ranking, amongst all actions, of the probability that the first T1 occurs within specified time bins before (left) and after (right) T2 triggers across learning. **i**, Quantification of pooled time bins from (**h)**. Repeated measures, 2-way ANOVA for learning stage vs. rank change. First T1 Before and After T2 Trigger groups differ across learning stage and total T1 probability rank change. (Proximal bins (0.3-1.8 sec): F(3,36) = 3.126. p=0.0376. Distal bins (2.1 to 3.6 sec): F(3,36) = 7.701. p<0.001). Post-hoc 2-tailed, Šidák multiple comparisons tests of all learning stage values compared to Starting Point values. **j,** Models for learning T1➔T2 sequences differing in initial time separations. Time not drawn to scale. **** p < 0.0001. *** p < 0.001. ** p < 0.01. * p < 0.05. n.s. – not significant. All bar and line plots indicate mean +/- S.E.M (**f-i**). See Supplementary Information for statistical/sample details.

The increased differential refinement favoring the proximal T2 could reflect increased refinement of T2 or reflect reduced refinement of distal T1 without refinement for T2. To distinguish between these interpretations, we analyzed changes in T1-T2 refinement curves relative to ‘Starting Points’ at which the refinement indices of T1 and T2 are most similar or are biased towards T1 rather than T2 (Methods). Slow Learners initially showed differential refinement favoring T2 from these Starting Points, and after reaching a maximum differential refinement favoring T2, called Turning Point, refinement begins to turn towards favoring T1 (Fig. 5c). By Turning Points, the median intervals of T1➔T2, but not T2➔T1 events decreased significantly relative to initial values (Fig. 5d, Extended Data Fig. 13c). Therefore, the median T1➔T2 interval decrease occurred before a decrease in the interval to perform the next sequence (T2➔T1) (Fig. 5e). Using these learning landmarks, we asked more rigorously how animals homed in on T1 vs T2 over time (Fig. 5f-g). Animals initially refined the action proximal to stimulation (T2), whereas T1 refinement occurred later, after Turning point (Fig. 5f-g). In contrast, Fast Learners show relatively little differential refinement over learning (Fig. 5c; Extended Data Fig. 14b-c). Finer analyses revealed that T1 was increasingly likely within seconds preceding T2 reinforcement events by Turning Point (Methods; Fig. 5h-i), even though T1 refinement was not yet apparent (Fig. 5f). After Turning Point, T1 refinement and increased sequence performance coincide with T1 becoming significantly more probable within seconds following T2 reinforcement events (Fig. 5h-i), indicating increased sequence re-initiation. These results demonstrate how animals can assign credit to sequences of temporally distant target actions that lead to reinforcement, following retrospective dynamics predicted by single action credit assignment. Specifically, actions most proximal to reinforcement are refined early on and the actions more distal to reinforcement become refined later, when they probabilistically start to occur within a few seconds of DA release.

## Discussion

Our results show that DA promotes credit assignment to specific actions and action sequences from a naïve state through a dynamic process whereby the entire behavioral repertoire is restructured and refined. During initial reinforcements there is rapid increase in frequency of not only of the target action, but also of actions in the repertoire that were similar to target. Dissimilar actions, however, decrease in frequency. This rapid restructuring of the entire behavioral repertoire based on similarity to target facilitates the credit assignment process. There is also an increase in actions that occur within a precise time window of a few seconds before and during, but not after, VTA DA neuron stimulation. With repeated reinforcement, gradual refinement unfolds to home in on the action that produces DA release. In the case of action sequences, both target actions in the sequence gradually become credited relative to their most similar actions. However, there is an interaction between the dynamics of refinement of the different target actions in the sequence and the temporal proximity to DA release. When sequences naturally varying in temporal separations between the two targets were reinforced, sequences with naturally short temporal distance between the two targets tend to refine together. However, credit assignment for sequences with naturally long temporal distances between the two targets is accomplished by earlier refinement for the actions most temporally proximal to reinforcement, followed by later refinement for the more temporally distal actions.

Previous synaptic and cellular studies^12, 13^ proposed that DA reinforcement may act retrospectively to reinforce behavior. By utilizing the closed loop system, we rigorously tested this prediction. Since retrospective reinforcement of behavior is not confined to the target action alone, it facilitates credit assignment to a stimulation-producing action even when reinforcement is delayed; stimulation-producing action pairs that tend to be performed closed together in time were learned much faster than pairs that tended to be performed far apart in time. Intriguingly, animals eventually learned to assign credit to distal stimulation-producing actions even in the latter scenario. This is characterized by a gradual process whereby early on, the median time interval between distal and proximal target actions decreased and the repertoire proximal to reinforcement was preferentially refined. As the distal target became significantly more likely to occur within second timescale distance prior to reinforcement, retrospective reinforcement of the correct stimulation-producing sequences became increasingly likely, resulting in whole behavioral refinement for the distal target as well, hence increasing sequence performance (Fig. 5j). This study has caveats that should be mentioned. The behavioral repertoire of the mouse in the open field is limited compared to more complex, naturalistic conditions. Furthermore, although our stimulation parameters produced similar dopamine release to that triggered by unsignalled sucrose consumption, these are not identical conditions. Therefore, the complexity of behavioral repertoire, and the frequency, duration and exact placement of dopamine activation^43^ could affect the exact window and rate of refinement/reinforcement. Still, the revealed principles of reinforcement and refinement during credit assignment should be generalizable.

Retrospective reinforcement of behavior^4, 19^ is thought to be mediated by DA modulation of an eligibility trace left by action potential-triggered synaptic plasticity^10^. Studies of DA action at the striatal synaptic level^12, 13^ indicate that retrospective reinforcement may occur on the order of a few seconds, but the behavioral consequences have remained elusive until now. Our behavioral findings agree with cellular studies that behavior occurring within a few seconds leading into DA stimulation are reinforced. Furthermore, they reveal an interaction between refinement process and temporal proximity to DA release –refinement becomes more pronounced if target actions occur within a few seconds before stimulation. The cutoff of retrospective reinforcement and refinement by phasic DA activities could explain the increase in sessions required to reach criterion frequency amongst animals that were reinforced for action pairs with initially longer median time separations. Similar actions have more similar and overlapping striatal neural ensemble activities^21^. Arrival of DA upon activation of action-specific ensembles may reinforce not only a specific action, but also similar actions. As striatal ensembles specific to actions are activated and a trial of eligibility traces is left temporally, DA arrival could mediate retrospective reinforcement of a spatially graded repertoire of actions within a few seconds, resulting in the observed behavioral learning patterns. Future studies would clarify how synaptic plasticity and cellular ensemble activities integrate to produce a dynamic refinement process, resulting in the behavioral principles for credit assignment revealed here.

## Methods

### Animals

All experiments were approved by the Portuguese DGAV and Champalimaud Centre for the Unknown Ethical Committee and performed in accordance with European guidelines. They were also performed according to National Institutes of Health (NIH) guidelines and approved by the Institutional Animal Care and Use Committee of Columbia University. 3-5 months old DAT-Cre male mice (*Mus musculus*) in the C57/BL6J background (RRID:IMSR_JAX:006660)^23^ were used. Animals were hetereozygous for the transgene and generated from a cross between heterozygote DAT-Cre males with C57/BL6J females. Animals were housed in standard light cycles (8am-8pm light, 8pm-8am dark).

### Sample Sizes, randomization, and blinding

For sample size, we applied a power of 0.8, significance of p<0.05, and standard variation of 20% of the mean. We determined sample sizes of 4-8 mice per group for different mean-based tests (matched pairs, 2 groups). No formal method of randomization was used; littermates were equally divided among the groups being compared. The experimenter was not blinded of the experimental groups. Optogenetic manipulations were performed automatically via a computer algorithm and not manually by the experimenter.

### Recombinant adeno-associated viral vectors, stereotaxic injections, and implants

750 nl of rAAV.EF1a.DIO.hChR2(H134R).eYFP or rAAV.EF1a.DIO.eYFP (3-4 x 10^12 vg/ml, AAV5, University of North Carolina Vector Core; 1-2 x 10^13 vg/ml, AAV1, Addgene, 27056-AAV1 and 20298-AAV1) were injected into each hemisphere of the VTA of 3–4-month-old DAT-Cre mice. For viral injections, the coordinates are AP - 3.52 mm, ML - +/- 0.35 mm, DV – 4.3 mm. Injections were made at 0.2 Hz pulses. Each pulse injects 4.6 nl volume. Injected needles were kept in place in the injection site for ∼15 minutes before withdrawal. For each mouse, a dual optic fiber cannula (200/240 μm diameter, 6 mm length, 0.7 mm center-to-center FLT, 0.22 NA; Doric, DFC_200/240-0.22_6mm_DF0.7_FLT) was placed 200 μm above the injection site and fixed to the skull. Next, a 4-position receptacle connector (Harwin Inc., M52-5000445) was fixed anteriorly to the dual optic fiber cannula, with its posterior edge set at −0.6 mm. Skull implants are then fixed with dental cement. A 4-position connector (Harwin Inc., M52-040023V0445) with pins removed from one end was used to cap the receptacle connector.

For photometry experiments, 3–5-month-old DAT-Cre males were used. The conditions used for VTA injections and implants were as above. Additionally, 1 μl and 500 nl of AAV9-hSyn-GRAB-rDA1m (2 x 10^13 vg/ml; Addgene, 140556-AAV9) were injected into the dorsal striatum (AP 0.5 mm, ML +2.1 (right), DV 2.3 (from brain surface)) and ventral striatum (AP 1.15mm, ML +1.65 (right), DV 4.2 (from Bregma)) , respectively. For photometry fiber implants, mono fiberoptic cannula were used (400/430 μm diameter, 4 mm length (dorsal striatum) and 6 mm length (ventral striatum), 0.37 NA, 1.25 diameter ferrule, flat; Doric, MFC_400/430-0.37_6mm_MF1.25_FLT (ventral striatum) and MFC_400/430-0.37_4mm_MF1.25_FLT (dorsal striatum)). Implants were inserted at a 22 degrees angle. For dorsal striatum implantation, the cannula entered the skull at AP 0.5 mm and ML 3.03 mm at 22-degree angle. The angled implant penetrated the brain from its surface for 1.92 mm. For ventral striatum implantation, the cannula entered the skull at AP 2.85 mm at 22 degrees angle, ML 1.65 mm. The angled implant penetrated the brain from its surface for 4.25 mm.

### WEAR motion sensor system

The WEAR motion sensor family was developed by the Champalimaud Hardware platform and Costa lab as a wired or wireless solution to obtain self-centered 9-axis motion data based on 3-axis accelerometer, gyroscope, and magnetometer (https://www.cf-hw.org/harp/wear). The wired version is a very small and extremely lightweight device (200mg) that can sample motion data up to 500 Hz and at the same time provide current up to 500mA that can be used to power LEDs for optogenetic experiments or stimulating electrodes. The wireless version is small and lightweight (∼1.8g) and can sample motion data up to 200 Hz while having the ability to provide up to 50 mA that can be used to power LEDs for optogenetic experiments or stimulating electrodes. The battery of the wireless WEAR allows recordings up to 4 h at 200 Hz sampling rate and even more at lower sampling rates. These devices communicate with the computers through a base station based on the HARP design developed by the Champalimaud Hardware Platform, which can be accessed through a software GUI to easily change sensor parameters to best fit the experimental needs. The base stations have several important hardware features such as 2 digital inputs and outputs, an analog input, 2 outputs for camera triggering, and a clock sync input and output that provides hardware-based synchronization. The sensor can be started or stopped by software or pin. The WEAR motion sensor family and base station are all open source (repository at https://bitbucket.org/fchampalimaud/workspace/projects/HP). Moreover, the WEAR devices are compatible with the Bonsai visual reactive programming software (https://bonsai-rx.org/), also open source, and allow the integration and synchronization of the streams of data being collected using the WEAR sensor with other data sources such as cameras.

Taking these specs and features together, the WEAR allows researchers to acquire high-resolution motion data wirelessly and for long periods of time, without being computationally very demanding. The 9-dimensional motion data acquired through WEAR is simple to process, easy to connect to analysis software, which allowed the fast online behavior classification that was fundamental for the experiments described in this paper.

### Open field experiment

One-month post-surgery, mice were habituated to head-mounted equipment over 2 days. On day 1, an actual or mock wireless inertial sensor (∼2.5 cm H x 1 cm L x 0.5 cm W with ∼ 2.5-3.0 cm antennae, ∼1.8 g weight) glued to the 4-position connector (Harwin Inc., M52-040023V0445) was attached to the implanted receptacle connector on the skull cap. Individual mice roamed freely in the home cage for 1 hour. On day 2, an actual wireless inertial sensor and mono fiberoptic patchcord (200/220 μm diameter, 0.22 NA; Doric DFP_200/220/900-0.22_2m_DF0.7-2FC) was attached to the skull cap via a mating sleeve. Patchcords were attached to 1x2 fiber-optic rotary joint (intensity division, 0.22 NA; Doric, FRJ_1x2i_FC-2FC) and mice roam freely in home cage for 1 hour. On open field recording day, sensor/patchcord habituated mice were anesthetized by isoflurane, attached to equipment, subjected to calibration protocol described below, and individually placed in an open field box inside a sound insulated chamber. The open field box is made of 410 x 400 mm grey opaque acrylic walls and a 410 x 400 mm white matte acrylic base. Individual mice were allowed to behave freely inside the box for 75 minutes. The wireless inertial sensor (∼1.8 g in weight, WEAR wireless sensor v1.1; Champalimaud Scientific Hardware Platform) conveys motion information sampled at 200 hz (set on WEAR v1.3.2 software; Champalimaud Scientific Hardware Platform) to a receiver base-station (Harp basestation v1.1 or v. 1.2, Assembly v0, Harp v1.4, Firmware v1.5; Champalimaud Scientific Hardware Platform), which conveys the information to the experimental computer running a Bonsai script (Bonsai^45^ editor v2.3.1, RRID:SCR_017218) to capture and record motion data and video information. Video was captured with a camera (Flea3 FL3-U3-I3Y3M(17450451), Point Grey Research) coupled to a 1/2” format lens (NMV-6WA, Navitar).

### Calibration

To ensure sensor stability within sessions, several approaches were employed. First, a coated mating sleeve was attached to the dual optic fiber cannula that sits immediately posterior to the sensor. The sleeve was thickened with black tape to a desired outer diameter such that it stabilized the sensor in the anterior-posterior direction. Second, the metal pins in the 4-position connector glued to the sensor were thickened with solder to stabilize their fit inside the receptacle connector in the skull cap. This protects against displacement in all directions. Third, stretchable black tape was wound around the base of the attached sensor and sleeve-covered cannula, further protecting against shifts in sensor positioning.

To control for possible variation in sensor positioning across sessions, a calibration approach was developed. Wireless inertial sensor was attached to individual isoflurane-anesthetized mice and the sensor was secured with the above strategies. Next, individual mice was placed in a custom-made calibration rig. The essential element of the rig is a vertical stainless-steel pole suspended above a stably secured table. In the setup used, the vertical pole was fixed to the horizontal edge of a vertically reversed “L” shape, stainless steel post assembly mounted on a breadboard (Thorlabs). The space between the lower end of the vertical pole and the table is enough for an individual mouse to slide underneath. The lower end of the vertical pole is fixed to a custom-made connector that resembles the connecting end of the fiberoptic patchcord. To perform calibration, individual isoflurane-anesthetized mice was securely attached to the vertical pole via a mating sleeve bridging the connection to the mouse’s cannula implant. Next, replicate readings of the immobilized inertial sensor were made on Bonsai. Next, mice were attached to the experimental patchcord and allowed to recover in home cage for 20 minutes or until individual mice are clearly recovered and behaviorally active. Individual mice were then placed in open-field box for experimentation.

Calibration involves rotating all accelerometer and gyroscope readings from the inertial sensor by a rotation matrix such that the final gravitational field vector of the stationary sensor, when mounted on the mouse and fixed to the calibration rig, is in a universal frame of reference whereby there is zero vertical tilt. In other words, the only non-zero acceleration is on the universal z-axis (pointing down). To accomplish this, the accelerometer pitch and roll orientation angles of the fixed stationary accelerometer were determined and then applied to calculate the rotation matrix. The rotation matrix is multiplied by the sensor accelerometer and gyroscope readings to remove the stationary vertical tilt from the sensor. To account for possible drift in gyroscope baseline over time, a daily reading of stationary gyroscope baseline was made with a mock cement skull cap attached to the sensor just before the start of each experimental day. The baseline gyroscope readings were subtracted from all gyroscope values before the rotation matrix is applied to sensor data.

### Action Selection

After open field run in the grey-walled box, off-line behavioral clustering was performed on calibrated sensor data. To identify the natural action repertoire of individual mice, we quantified behavior using acceleration and gyroscope time series features in a similar fashion as described previously^21^. For the ground truth analysis, we used: 1.) Gravitational acceleration (GA) along the anterior-posterior (A-P) axis for the discrimination of postural changes - GAap. 2.) Raw sensor acceleration along the dorsal-ventral (D-V) axis to quantify movement momentum – ACCdv. 3.) D-V axis of gyroscope to extract head head-body rotational information – GYRdv. 4.) Total body acceleration to differentiate resting state from movement.

Total body acceleration (TotBA) was defined as:

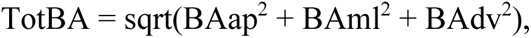

where BAap, ml and dv represent the body acceleration of the anterior-posterior, medio-lateral and dorsal-ventral axis, respectively. We calculated each individual BA component by median-filtering the raw acceleration signals followed by a fourth-order Butterworth high-pass (0.5Hz) filter. For the gravitational acceleration (GA) axis, the BA components were subtracted from the median filtered raw signal axis.

All four time series features were binned into non overlapping 300 ms long window segments^26^. The values of each bin and per feature were then discretized, using fixed thresholds, producing a summary distribution of each segment. For GAap and ACCdv we used 10 equal size threshold values, plus two added bins between the limits and infinity to capture an approximated distribution of values within each window bin. For GYRdv we used 5 thresholds (0, ±50, ±100) to discriminate left and right turns. For TotBA, a single threshold was used to separate moving from resting. The threshold was kept constant for all experiments and was set to the average value separating the bimodal distribution of logTotBA (natural logarithm of TotBA feature). For each 300-ms window segment we get four resulting histograms, one for each feature. The feature histograms were individually normalized to obtain probability distributions and used to calculate the pairwise similarities between segments.

We used the “earth mover’s” (EM) distance as a measure of similarity^25^:

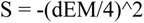

where dEM is the sum of the normalized EM distances for the 4 features (GAap, ACCdv, GYRdv and TotBA) defined above. The bin normalizations constrain S values within the range [-1,0], specifically, -1 and 0 define the maximum dissimilarity and identity between the two probability distributions, respectively. Finally, to produce a continuous unbiased classification of behavioral states, the similarity measures were clustered using affinity propagation^20^, with the preference parameter set to the minimal value of the similarity matrix; this particular value was used for its stable number of behavioral clusters within its range.

Using the behavioral clusters identified by affinity propagation clustering of the grey open field behavior^13^ as a ground truth for the true identity of each 300 ms histogram, we were able to simulate and evaluate the precision with which the Earth Mover’s Distance (EMD) metric^21, 25^ could be applied for cluster matching online. Notable difference between the EMD metric used here is the use of the 4 features mentioned above rather than the 3 features used previously^21^, as well as the multiplication of the similarity score by -1 such that the range of possible scores from maximal identity to dissimilarity is 0 to 1, respectively. Although the EMD cluster matching outcome correlates strongly with affinity propagation clustering, some false positive and false negatives may occur. Several filters were set to optimize cluster selection for reinforcement: 1.) We selected for clusters that show low false positive rate (<5.5%) and below the 60^th^ percentile false positive rate amongst all clusters per animal. 2.) We selected against clusters with high false negative rates (> 90^th^ percentile of clusters per animal). 3.) We selected against clusters that tend to be performed serially within a short time interval. We calculated the probability that a target cluster or its top 5 most similar clusters (determined by EMD score) would reappear 3-18 seconds after the first occurrence of the target cluster. Clusters that tend to be repeated either by itself or have a high probability of having similar clusters appear within this 15 second window (> 90^th^ percentile for median and range of probabilities of cluster appearing in window) were removed from selection pool. 4.) We filtered against clusters whose matching by EMD would be more sensitive to anterior-posterior shifts of the inertial sensor (although we already protected against this possibility with the safeguards above) (> 90^th^ percentile for percent deviation from original cluster matching after shifts of accelerometer reading in the anterior or posterior direction). For each cluster, percent deviation is calculated first by summing up the total absolute cluster matching changes from original cluster matching data in the anterior and posterior shifted datasets. Next, the sum of deviation in the two altered datasets is divided by two and then divided by the total of cluster calls from the original dataset, and multiplied by 100 to get percent deviation from original cluster matching result. 5.) We selected for clusters that show fully accelerating movement (cluster exemplar value of 1 (maximum value) in the body acceleration feature bin of histogram). We did not choose the exact same type of actions across animals to improve generalizability of discovered learning principles. However, action types comprise a mix of complex behavior visually similar to those described previously^21^. To choose dissimilar clusters per animal, an algorithm was written filtering clusters of each animal’s repertoire based on the feature histogram values of each cluster’s representative, or exemplar. Thresholds were set along the GAap and GYRdv features to divide cluster exemplars based on the distribution of values within these feature histograms. For each repertoire, all histogram values from all cluster exemplars are pooled to create a pooled histogram. The range of bins with non-zero values for each feature are identified. The algorithm then filters cluster exemplars in the repertoire for non-zero values in the high, medium, low, or high+low value bins. For example, action A identification occurs by selecting for a cluster exemplar with median counts falling in the high GAap and GYRdy value bins. action B would then be selected by filtering for an exemplar with median counts falling in the low GAap and GYRdy value bins. This process was alternated from animal to animal, such that the selection criterion for Action A and B would continually be reversed (ex. Animal 1’s Action A (high GAap / GYRdy values) and Action B (low GAap / GYRdy values) ➔ Animal 2’s Action A (low GAap / GYRdy values) and Action B (high GAap / GYRdy values) ➔ Animal 3’s Action A (low GAap / GYRdy values) and Action B (high GAap / GYRdy values) ➔…). This results in action pairs that are highly dissimilar within animals and Actions A and B that are broadly distributed across the action space. Dissimilar behaviors differ in complex ways, but visually can be roughly described as varying in some example dimensions such as vertical vs. horizontal posture, heads down vs. heads up and left vs. right turns, etc. EMD similarity scores comparing action A to action B almost always, except for 1 ChR2-YFP animal, fall in the more dissimilar end of a distribution of scores created by comparing action A to all actions in each animal. Hereafter, clusters will be referred to as actions.

### Closed-Loop Optogenetics

For close loop optogenetics, a computer running a Bonsai script captured and recorded wireless sensor motion data and video information as described above in grey-walled open-field experiment. Here, data is also streamed to a custom MATLAB code which analyzes action composition changes over the course of action reinforcement, we used the EMD metric^21^ to label individual 300 ms motion histograms with an action ID. For each arriving 300-ms segment we calculate the EMD distance between each cluster exemplar (or representative) of the ground truth cluster library from the grey open field behavior recording. The motion features histogram is assigned to the action for which comparison with the exemplar gave the lowest EMD score (most similar to target) amongst all comparisons. Decision making for stimulation has a range of 35-55 ms time gap between action performance and sent decision for stimulation. To trigger optogenetics, a Multi-Pulse Width Modulation (PWM) generator (Harp Multi-PWM Generator hardware v1.1, Assembly v1, Harp v1.4, Firmware v1.1; Harp Multi-PWM Generator software v2.1.0; Champalimaud Scientific Platform) converts each decision to trigger laser into electrical signals for 15 light pulses of 10 ms pulse duration at 25 Hz, with each train of pulses occurring over 600 ms and at 25% duty cycle. The multi-PWM signal is passed through a 12 V, 7.2 W amplifier (Champalimaud Scientific Platform) and fixed frequency driver (Opto-electronic, MODA110-D4-30 (2001.320220)) to control the activities of a 473 nm, blue low noise laser (Shanghai Dream Lasers Technology, Co, Ltd. SDL-473-200T), which was sent through an acousto-optic modulator (Opto-electronic, MTS110-A3-V1S (1001 / 330433)). The laser component that is modulated is then reflected by a mirror and funneled to a mono fiberoptic patchcord, which is then coupled to a commutator. The output laser is then passed through a dual-optic fiber patchcord and connected to the implant cannula. Power adjustment out of the tip of patchcord was made so that ∼5mW was emitted from each end of the dual optic fiber cannula. To ensure common time stamps from different channels, a clock synchronization device (Harp Clock Sync v1.0; Champalimaud Scientific Platform) was performed between the basestation and multi-PWM device.

### Single action sequence reinforcement

Mice were placed in a white open field box for closed loop reinforcement protocol. A white open field was used instead of the earlier grey open field to minimize habituation effects, which would lead to reduced initial spontaneous behavior during closed loop protocol. Individual mice were subjected to a single session of protocol each day, with sessions following each other on consecutive days. The white open field box is made of 410 x 400 mm white matte acrylic walls and a 410 x 400 mm white matte acrylic base. To acquire baseline behavior, individual mice were allowed to behave freely inside the box for 30 minutes on the first action A reinforcement session. Closed loop reinforcement by blue laser stimulation of VTA DA neurons were made available for 60 minutes. 90 minutes of closed loop reinforcement were made available for individual mice during sessions 2 and 3. For session 4, an extinction protocol was carried out comprising of 20-minute maintenance of reinforced behavior with laser availability, followed by 60 minutes of extinction of reinforced behavior without laser availability, followed by 20-minute re-acquisition of reinforced behavior with laser availability. To select for action B, a repeat of the protocol described above for action A was performed starting on the day following extinction protocol of action A. Upon completion of the reinforcement and extinction protocols for action B, a contingency degradation protocol was performed comprising of 20-minute maintenance of action B with laser availability, followed by 60 minutes of contingency degradation of reinforced behavior by triggering laser randomly, followed by 40-minute re-acquisition of reinforced behavior with laser availability for action B performance.

### Photometry experiment

One-month post-surgery, mice were habituated to head-mounted equipment for 2 days. On day 1, habituation was made to wireless inertial sensor as described above. On day 2, a multi-fiber bundled patch cord (3 fiber bundle, 400/440 μm diameter for a maximum of inner diameter at 900 μm, 0.37 NA, 3.5 m long, 1.25 mm fiber tip diameter, low-autofluorescence; Doric, BBP(3)_400/440/900-0.37_3.5m_FCM-3xMF1.25_LAF) was attached to individual mice in addition to the wireless sensor and optogenetic patchcord. Individual mice were allowed to habituate to the equipment for 1 hour in its home cage. On photometry recording day, mice were subjected to 30 frames per second photometry recording (FP3002, Neurophotometrics), with 75-150 μW 560 nm LED illuminating rDA1m, and equivalent closed loop optogenetic parameters described above were used. To test for DA release in the context of closed loop optogenetic setup, an average of 30 hits of blue light were delivered randomly within the span of 30 minutes. To evaluate DA release in the context of food reward, mice were placed on food deprivation protocol and kept within 85% of original weight. Mice were placed in an operant chamber with a nosepoke linked to a lick detector (PyControl). Each lick detection triggers dispensing 2 μl 10% sucrose. Since animals tend to accidentally trigger lick detector at the beginning of sessions, between 40-50 sucrose dispensing events were gathered per animal and rDA1m activities associated with the last 35 rewards of the session were used for analysis.

### Two action sequence Reinforcement

Two action sequence reinforcement occurs as follows: after sensor/patchcord habituation and grey open field behavior recording, offline behavioral clustering and action filtering were performed as for single action reinforcement. For each animal, median time intervals between all possible pairs of actions during open field were calculated as described above. Across animals, T1/T2 pairs with median T1➔T2 interval values varying between 2 and 10 seconds were selected. To control for variation due to order of movement orientations, various combinations of target action T1 having low GAap / directional GYRdy values to target T2 having high GAap value / opposite directional GYRdy values, were chosen for reinforcement.

On the first reinforcement session, a 30-minute baseline was taken when laser stimulation was not available for reinforcement. Laser became available for reinforcement in all subsequent sessions until extinction experiment. During reinforcement periods, when closed-loop system detects performance of the proximal action (T1) of interest, the algorithm enters a state where laser is triggered upon performance of the distal action (T2), regardless of the amount of time that has elapsed between the latest T1 and T2. On Session 1, 60 minutes of laser availability was given while in all subsequent reinforcement sessions, 90 minutes of laser availability was given.

### Histology and Immunohistochemistry

After behavioral sessions were completed, mice were deeply anesthetized with isoflurane and perfused transcardially in PBS and then 4% PFA/PBS. Dissected brains with skulls attached were perfused in 4% PFA in PBS at 4 degrees Celsius overnight. The next day, brains were rinsed 3 times in PBS. Next, brain regions including VTA and implants were sectioned by vibratome into 50 or 100 μm slices. Slices are then subjected to immunohistochemistry using the reagents below. Standard immunohistochemistry protocols were applied to stain for the following reagents - Rabbit anti-GFP 488 conjugate (1:1000; Molecular Probes A21311). Mouse Anti-TH (1:5000; Immunostar Th 22941) with Goat Anti-Mouse - IgG (H+L) Highly cross-adsorbed secondary antibody - Alexa Fluor647 (1:1000; ThermoFisher, A-21236), DAPI (1:1000 of 20 mg/mL stock; Sigma, D9542).

### Imaging

Zeiss Axio Imager M2 microscope was used to acquire brain section pictures. 10x tiled images were taken through the relevant fluorescent channels. The M2 is equipped with a fast Colibri.7 LED illumination for excitation of fluorophores. Images are captured with a high-sensitivity monochromatic sCMOS camera (Hamamatsu Orca Flash 4.0 v2). The objective used for the images is a ZEISS Plan-ApoChromat 10x/0.45, which allows to resolve up to 577 nm when using a wavelength of observation of 520nm and it is fully corrected for chromatic and spherical aberrations. Implant locations were determined using standard mouse atlas^46^.

### Single action reinforcement analyses

For target action frequency analysis, we analyzed frequencies within 25-minute windows at 4 time points: Baseline (before first reinforcement trigger), Early (after first reinforcement trigger in Session 1 (action A) or 5 (action B)), Mid (after 2-minute mark in Session 2 (action A) or 6 (action B)), Late (after 2-minute mark in Session 3 (action A) or 7 (action B)). For 3D action repertoire plots, baseline normalized frequencies were plotted and actions whose time series include NaN or Infinity values were discarded from the plot. (Plotted actions: 509 of 514 actions, 15 ChR2YFP animals (action A); 427 of 443 actions, 13 ChR2YFP animals (action B); 355 of 356 actions, 10 YFP animals (action A); 341 of 356 actions, 10 YFP animals (action B)).

Three parameters were assessed for rapid behavioral adaptation following cumulative closed loop reinforcements: latency between Target A triggered reinforcements, Target A frequency and average behavioral similarity to Target A. Latency refers to time interval between consecutive triggers. To calculate the latency parameter, the average latency between 10 consecutive Target A triggered reinforcements following a specified number of cumulative reinforcements were taken and then normalized by the average latency taken over the final 10 baseline Target A instances that in simulations would have triggered reinforcement. To calculate the frequency parameter, the frequency of Target A triggered reinforcements over the course of 1 minute following a specified number of cumulated reinforcements were taken and then normalized by frequency of the final 10 baseline Target A instances that in simulations would have triggered reinforcement. To calculate the behavioral similarity parameter, the average behavioral similarity (EMD score) to Target A between 10 consecutive Target A triggered reinforcement events following a specified number of cumulated reinforcements were taken and then normalized by the corresponding value taken over the final 10 baseline Target A instances that in simulations would have triggered reinforcement.

### rDA1m Fiber Photometry Analyses

To evaluate DA release in the context of food reward, the delta F/Fo signal was plotted for rDA1m signal aligned to lick detection/reward trigger (Fig. 1f). The baseline Fo value was taken as the median rDA1m raw fluorescence signal of the 10 time points (333.33 milliseconds) preceding the trigger event. To test whether DA release is triggered in the context of the closed loop system, the activity of the rDA1m sensor was quantified. Delta F/Fo was calculated by subtracting baseline value from each fluorescent rDA1m value of a smoothened time series (smooth function, default moving average filter, MATLAB), and then dividing the outcome by the baseline value. To account for control ChR2-independent effects, the average delta F/Fo trace of ChR2-YFP animals were subtracted from the corresponding average trace of YFP animals, giving the differential delta F/Fo used for the plots. The standard deviation of ChR2-YFP minus YFP curves were obtained by taking the square root of the sum of squared variances of ChR2-YFP and YFP delta F/Fo curves.

### Categorizing behavioral actions by temporal dynamics

To categorize behavioral actions by temporal dynamics (Fig. 1m, Extended Data Fig. 6), moving mean of action counts was used as input. Various window sizes were examined; 2.5-minute windows moving at 300 ms steps were found suitable for analyses. The baseline frequency (f0) was the average of 5 minutes of moving mean data preceding the first reinforcement event. Early frequency rate (f1) was the average of 30 minutes moving means immediately following the first reinforcement event. Mid- and Late frequency rates were taken from Day 2 (f2) and Day3 (f3), respectively. f2 and f3 rates were calculated from the beginning 30 minutes period after moving windows has accumulated enough bins (2.5 minutes) following the start of the session. Significant positive modulation above baseline was judged if in 500 consecutive moving windows (2.5 minutes period) in Early/Mid or Late stages the frequency rate of all bins were greater than the 99^th^ percentile bin of baseline frequency. Significant negative modulation below baseline was judged if in 500 consecutive moving windows (2.5-minute period) in Early/Mid or Late stages the frequency rate of all bins were less than or equal to the 5^th^ percentile bin of baseline frequency. Actions that showed both significantly positive and negative modulation at Early/Mid or Late stages when compared to baseline were delegated to positive modulation group. For figure plotting, time-course median frequencies of action dynamic types were downsampled 10-fold (Extended Data Fig. 7a). To examine the distribution of action dynamic type frequencies in terms of target similarity, a binning by raw EMD score (0.5 score binwidth) was used because this allowed for clear visualization of the relationship between target similarity and frequency (Extended Data Fig. 7b). Alternatively, percentile binning of EMD score was also used and gave similar trends (Extended Data Fig. 7c).

### Criterion for action dynamic types

Action dynamics were grouped according as follows: 1.) Increasing actions showed significant increase in f0 to f1/2 and f1 to f2/3 comparisons and showed either significant increase or unchanged frequency in f1/2 to f3 comparisons. 2.) Sustained actions showed significant increase in f0 to f1/2 comparisons, and unchanged frequency in f1 to f2/3 and f1/2 to f3 comparisons. 3.) Transient actions showed significant increase in f0 to f1/2 comparisons, and significant decrease in f1/2 to f3 comparisons. 4.) Decreasing actions showed significant decrease in f0 to f1/2 and f0 to f3 comparisons. 5.) Other actions were all remaining actions that did not fall in the above groups. In Extended Data Fig. 7a, only dynamic subtypes with more than 10 members are shown.

### t-SNE and hypervolume analyses

Data from 5-minute portions of baseline (Session 1 pre-reinforcement portion), early (Session 1 reinforcement portion), mid (Session 2) and late (session 3) were pooled to calculate pairwise EMD similarities between all individual action instances, creating a similarity matrix. An individual action instance’s similarity distance against all other actions in the dataset specify the action’s position in behavioral space. The first 50 principal components of this dataset were embedded into a 2D behavioral space with t-SNE. Hypervolume analysis was performed on the using the dynamic range boxes method^34^, and implemented with the dynRB package in R (RRID:SCR_001905). To account for more information in n-dimensional space, the first 250 principal components of the similarity matrix (4,000 total action instances randomly sub-sampled in equal numbers from each baseline/reinforcement portions) were used to calculate hypervolume overlaps. The port parameter (proportion of a particular action cluster’s hypervolume overlapping with that of the target cluster hypervolume) was evaluated. To evaluate changes to transiently increased dynamic type percentage per animal after accounting for overlapping hypervolumes, all action clusters whose hypervolumes had non-zero overlap with the target hypervolume were removed to re-calculate the percentages of each dynamics type per animal. Data is presented in Extended Data 8.

### Extinction analyses

10 minutes portions from different time windows along the extinction protocols (Session 4 for action A and Session 8 for action B) were chosen (Extended Data Fig. 4c (action A), Extended Data Fig. 9g (action B). Early maintenance (M^1^) starts from the first instance of target action performance in the session. Late maintenance (M^2^) is the portion preceding the first performance of target upon extinction. Early extinction (E^1^) begins at the first instance of target performance upon extinction. Late extinction (E^3^) is the portion preceding the first performance of target upon re-acquisition. Mid extinction (E^2^) begins at the midpoint between the starts of E^1^ and E^3^. Early re-acquisition (R^1^) starts at the first performance of target upon re-acquisition condition. Late re-acquisition (R^2^) is the final portion of the extinction protocol.

### Action burstiness analysis

To evaluate action burstiness, or dispersion, we used Fano factor (variance/mean) as a measure (Extended Data Fig. 4f). A survey of moving mean frequencies of reinforced actions across animals suggest that actions are more dispersed during the extinction phase, but the timescale with which this may occur is variable. To identify a suitable timescale to detect dispersion across reinforced actions, we screened a range of window sizes (600 ms to 5 minutes windows in 600 ms steps) with which to calculate moving window frequencies, and then calculate Fano factor in varying time segments. We chose a moving window of 15 seconds (50 x 300 ms action units) to construct moving mean frequencies. This window size consistently gave decreased Fano factor in baseline vs. maintenance session across animals, a result that would be expected as reinforcement led to stable target action performance.

### Single action reinforcement, inter-target, and inter-action interval analyses

To quantity inter-target action intervals (Fig. 3a-b), the median amount of time that transpired between the start of successive target actions over the course of a time window was calculated. The time periods analyzed were: 1.) Baseline from the start of Day 1 (Sessions 1 and 5 for action A and B, respectively) until the first reinforcement event. 2-4.) Days 1 to 3 reinforcement. For reinforcement periods, behavior from the start of the first reinforcement event of that session until the end of session were analyzed. We considered the possibility that including the time interval between consecutive repeating of target actions (resulting in an inter-target action interval of 300 ms) would greatly affect the result. To test this, we removed values collected from consecutively repeating target actions. However, this did not affect result interpretations. Thus, we included intervals from consecutively repeating target actions in the presented analyses. For single action reinforcement, the median amount of time between the closest occurring action of interest and target action was calculated for both pre-target and post-target intervals.

### Multinomial logistic regression predicting action dynamic types

To test whether intrinsic and baseline action properties are predictive of classifiable action dynamics during single action reinforcement from naïve state, two factors were considered. The factors are Earth Mover’s Distance (EMD) similarity of action to target and median time interval of closest action of interest preceding target appearance at baseline condition (Fig. 3c-g).

To perform multinomial logistic regression, data from both dependent variables were normalized to z-scores. Transformed data were tested for collinearity by examining scatter plots, Pearson’s correlation coefficients, Variance Inflation Factors (VIF) and condition indices. The two variables showed some correlation, but the coefficient value was not above typical thresholds^47, 48^ and direct collinearity diagnostics did not show significant collinearity (Pearson’s correlation: 0.61 < 0.8^47^, VIFs: 1.6 < 5-10^49^, condition indices: 2.0< 10-30^50^). Multinomial logistic regression was performed using MATLAB functions mnrfit and mnrval. The mnrfit function uses the iteratively weighted least squares algorithm to find the maximum likelihood estimate of the coefficients in a multinomial logit model. In such a model, the relative risk of being in one action dynamic type category versus the reference group (Decreased dynamics type) is expressed as a linear combination of predictor variables, each with its own beta-coefficient. The mnrval function predicts the category probabilities. Non-Target A actions from all animals from reinforcement of action A were included except those whose reinforcement dynamics were previously classified as “Other” types (n = 30 actions from a total of 514 actions, 15 ChR2-YFP animals). Model accuracies were assessed using a 20-repeat, 10-fold cross-validation approach for a total of 200 unique models for Real data, and 10,000 unique models from 50 shuffled datasets.

To evaluate multinomial logistic regression, the deviance measure was used to judge model fitting (Fig. 3f). The deviance of the fit is a goodness-of-fit statistic that is calculated as twice the difference between the maximum achievable log likelihood (in a saturated model where data fits perfectly) and the corresponding likelihood in the fitted model (the actual model of interest). Model performances were judged by area under precision-recall curve as this criterion is suitable for imbalanced categories in the data^38^ (Fig. 3g). A model containing both dependent variables was found to outperform that of any single variable in data fitting, even after consideration for penalties for an extra factor (Akaike Information Criterion; 2 multiplied by 2 independent variables (2k = 4) as penalty added to deviance of 2-factor model^51^). Further, the 2-factor model outperformed both 1-factor models in predicting true positives of Decreased type, while performing just as well as the Similarity factor model in predicting true positives of Sustain Increased and Transient types (see table in Supplementary Information). The 2-factor model outperformed the Time preceding Target factor model in predicting all types (see Fig. 3f(III) tab in Supplementary Table). The lack of significant collinearity between dependent variables was supported by the stability of two relevant parameters, beta-coefficient directions and significant p-values, across 200 cross-validation models and single- and double-factor regression conditions (See Supplementary Information for tables).

### Dopamine retrospective window analysis

To test whether reinforcement was selecting for behavior before, during or after stimulation, we originally tracked how initial baseline distributions of single actions surrounding stimulation relate to single action frequencies upon reinforcement. However, we were not able to tell whether single action frequency changes upon reinforcement were associated with each action’s baseline distribution before or after stimulation (the distribution was highly symmetric due to lack of contextualization).

To improve contextualization, we analyzed first order transitions. This provided enough context for us to distinguish between behavior occurring before, during or after stimulation. Thus, we were able to identify action transitions specifically associated with specific temporal positions relative to stimulation, track these action transitions as reinforcement commences, and attribute their frequency changes to their temporal positioning surrounding stimulation.

To analyze whether DA reinforces actions proximal to target, rates of action transitions occurring close to reinforced action were examined during baseline and over the course of closed loop reinforcement (Fig. 3h-j). First, 600 ms action transition events (ex. X ➔ Y) occurring from 2.4 seconds before to 2.4 seconds after each theoretical target-triggered laser stimulation (600 ms in length) during baseline condition were examined. Next, all possible transitions occurring within specific 600 ms sliding windows within the defined time range were counted for each animal. Next, the relative performance of each action transition type in a specific sliding window against all sliding windows was calculated by dividing a particular action transition type’s count within a specific sliding window by the total number of the same action transition type across sliding windows. This gives probabilities across sliding windows. Next, action transition probability within a sliding 1.2 second window (“within” region in Fig. 3h schema; containing a total of three action transitions) relative to surrounding temporal environment (“outside” region in Fig. 3h schema; 3.6 seconds) was derived by subtracting the probability of a particular action transitions type occurring inside an “outside” region from the probability of the same type occurring in the “within” region of interest. This will be called the differential probability. Next, action transition types that showed greater or equal to a threshold of 0.001 differential probability within sliding 1.2 second windows of interest over the corresponding surrounding “outside” windows were filtered and kept for the next step. This marks the selection for enriched action transitions. Next, for each sliding 1.2 second window, transition count data from above was analyzed to select for action transition types that occurred between 2 to 6 times during the 30 minutes baseline period (0.067 to 0.2 action transitions per minute). The count range was chosen to filter out single events while selecting for action transitions with low initial frequencies over the baseline period and analysis time range. Since the range of probabilities of specific action transition types could vary greatly between different sliding 1.2 second windows, filtering as above also balances the distribution of action transition probabilities amongst all action transition types analyzed across sliding 1.2 second windows. The above process results in a list of action transition types enriched for each sliding 1.2 second window, and baseline normalized frequencies of these action transition types upon reinforcement in subsequent sessions were calculated. Note that baseline normalized frequencies were calculated from all occurrences of specific action transition types, regardless of their time distance in relationship to target occurrence. Baseline normalized frequencies of individual action transition types were averaged within animals and the means between animals are averaged to produce animal-balanced results (Fig. 3i-j). Identical data trends and conclusions could be reached even if baseline normalized frequencies of all action transitions were used for analyses.

### Two action sequence experiment analyses

Two action sequence frequency was quantified in terms of laser triggers per minutes (Fig. 4b, d-g). To assess learning across animals, the baseline frequency was subtracted from frequencies of all reinforcement sessions (Fig. 4b, d). A criterion baseline subtracted frequency of 3.2 triggers per minute was set after considering the range of baseline subtracted frequencies observed in the open field and reinforcement sessions all animals. The criterion is set such that it is > 20 % above the highest baseline-subtracted frequency value seen at open field condition. The criterion point consistently falls above the open field frequencies of all animals and marks the rising phase of all reinforcement frequency curves.

T1➔T2 intervals were quantified as the time distance between the end of the latest distal action (T1) and the end of the proximal action (T2) that triggers laser. T2➔T1 intervals were quantified as the time distance between the end of T2 that triggers laser and the end of the next closest T1. To produce equivalent measures in open field and baseline conditions, laser trigger events were simulated by scanning across the data as if reinforcement was available.

Significance testing was performed on 14 of 15 ChR2-YFP animals that reached criterion frequency (ChR2-YFP Criterion) (Fig. 4e). The lone animal that did not reach criterion frequency was removed because the T1➔T2 median interval was still very high after session 10. This animal was subsequently subjected to single action reinforcement protocol to assess its ability to learn T1 and subsequently T2 (Extended Data Fig. 10a-b). Next, the animal was again subjected to T1➔T2 reinforcement protocol (Extended Data Fig. 10c). These results indicate that this animal was capable of action learning for both T1 and T2 separately, and for T1➔T2 sequence after learning of each individual action.

Reinforcement sessions for the 14 ChR2-YFP animals that reached beyond criterion frequency continued until the T1➔T2 interval has been decreased to below at least a median of 2 seconds (Fig. 4h). As YFP animals do not decrease the T1➔T2 median interval over sessions, we stopped reinforcement at session 20.

### Two action sequence extinction

14 of 15 ChR2-YFP animals that passed criterion sequence frequency were subjected to the following extinction protocol. The remaining animal was subjected to an earlier extinction protocol spaced over two sessions and had a longer stretch of extinction time (Extended Data Fig. 10). This animal showed loss of sequence performance over the extinction period but became largely inactive by the reacquisition period, slowing reacquisition within the allocated time window. This led to shortening of the extinction protocol to 40 minutes of extinction and having all conditions within a single session as described here. Extinction session begins with a 25-minute maintenance period for two action-sequence reinforcement, followed by a 40-minute extinction period when laser was inactive, followed by a 25-minute re-acquisition period whereby reinforcement was made available again (Fig. 4f-g). To quantify performance for plotting, frequency was calculated over 5 minutes bins and then normalized to the last 5 minutes bin of the maintenance condition (Fig. 4f). For significant testing, raw frequencies were analyzed at the last 5 minutes of maintenance, extinction, and re-acquisition conditions (Fig. 4g).

### Two action sequence refinement

To measure refinement for T1 and T2 in the two-action sequence (Fig. 4k, Fig. 5), actions that were uniquely related to one but not the other were identified. Actions performed by each animal in their open field repertoires were ranked by their EMD similarity scores to T1 or T2. The top-12 actions (within action repertoires ranging between 30-40 actions) most similar to either T1 or T2 were identified. Actions common to both T1 and T2 in these lists were removed, leaving actions uniquely similar to T1 or T2. We required at least 3 non-target actions to be uniquely related to each of T1 and T2. One of the animals did not meet this requirement, because less than 3 actions were uniquely similar to each of T1 and T2 when considering the top-12 actions related to T1 or T2. For this animal, we relaxed the stringency by considering actions that uniquely belong as the top-9 actions most similar to either T1 or T2. We took the median target-normalized frequency of these uniquely similar actions to T1 or T2 as the refinement index. A refinement index of above or around 1 indicates little to no refinement of uniquely related actions to target. Refinement index below 1 indicates refinement relative to target; the lower the score the more refinement. Refinement curves were smoothened using the Savitzky-Golay filter to improve visualization of trends (Fig. 5a). To better compare the progress of refinement between T1- and T2-related actions, refinement indices were scaled such that the minimum value amongst all sessions for individual animals would be zero and target-normalized median frequency of 1 would remain at a scaled value of 1 (Fig. 5a, f).

### Relationship between T1➔T2 interval and sessions to criterion frequency

To describe the trend in a T1➔T2 interval vs. sessions to criterion frequency scatter plot, non-linear sigmoidal fit was tested against a 4^th^ order polynomial fit (Fig. 4l). A linear fit was also tested. Sigmoidal fitting gave the best result. The same fitting was tested for T2 ➔ T1 interval vs. sessions to criterion frequency, but the fit was poor, and midpoint was unstable. For the T1➔T2 sigmoidal curve, half-maximum was 2.59 sessions to criterion frequency and midpoint was 4.69 seconds of open field median interval. The half-maximum value was used to divide ChR2-YFP animals into slow (above half-max) and fast (below half-max) learners. Identical grouping of fast and slow learners could be obtained by taking the median value across animals as separation point.

To test whether reinforced action pairs differ in initial inter-trigger intervals (Extended Data Fig. 11d), we simulated and calculated the natural median interval occurring between T1➔T2 performances (called median inter-trigger (simulated) interval) and asked whether variation in this parameter at baseline predicts learning outcome (sessions to criterion frequency). To test whether the initial T1➔T2 median interval influences learning after matching for reinforcement numbers over learning (Extended Data Fig. 11e-g), learning was examined by calculating the average number of sequence performance per unit time, with the time range covering spans of 200 reinforcements. To further account for differing initial and eventual sequence frequency over time, we smoothed all individual learning curves with a Savitzky-Golay filter and then scaled the curves to the frequency in the initial 200 reinforcement bin (value = 0) and to the maximal frequency (value = 1). To ensure that a similar conclusion regarding the relationship between initial T1➔T2 intervals and learning could be derived, we plotted initial T1➔T2 median intervals against the number of reinforcements cumulated upon reaching criterion frequency (25% of maximal scaled frequency) and tested for ability to fit a sigmoidal curve as in Figure 4l.

### Differential refinement analyses

The difference in area between T1 and T2 scaled refinement curves over sessions was used to assess the relative refinement status between T1 and T2 over sequence learning (Fig. 5b). The difference in areas were summed up using the trapezoid method across sessions until the session when both T1 and T2 has or had reached minimal scaled refinement. Next, the relationship between open field median interval and average difference in area under T1 – T2 refinement curves per session was tested. A per-session metric was employed to control for variations in total sessions across animals. Linear regression proved most suitable for fitting (Goodness-of-fit: R2 = 0.66). The fit for T1➔T2 linear line was y = 0.1893x – 0.7050. Slope was significantly non-zero (p = 0.0004). The same fitting was tested for T2 ➔ T1 interval vs. difference in area under T1 – T2 refinement curves per session (y = 0.00736x + 0.1356), but the fit was poor, and goodness of fit was low (Goodness-of-fit: R2 = 0.07). The slope was not significantly non-zero (p = 0.7063).

It is possible that for fast learners, the differential refinement favoring proximal action (T2) is only seen in very early parts learning (ex. intra-session blocks within session 1 and 2). To account for this, we probed intra-session refinement dynamics for fast learners by dividing each session into three x 30-minute blocks and repeated the refinement analyses in Fig. 5b (Extended Data Fig. 13a). We performed the analysis only up to the session of criterion frequency (usually one of the early sessions for fast learners) to avoid diluting out the effect from later sessions when differential refinement tends to even out as animals learn the sequence. We further visualized differential refinement dynamics for fast learners similarly to Fig. 5c.

The intra-session analysis showed that fast learners underwent diverse differential refinement dynamics. Some fast learners show differential refinement within session for the proximal action (T2) while others show the opposite (favoring T1 early on). Some did not show clear preference for either. Importantly, these higher resolution results were consistent with the analyses done based on whole sessions data (Fig. 5b). A similar linear relationship between open field T1➔T2 median interval and differential refinement (change in area under the T1-T2 refinement curves (per time block)) was observed (R-squared is 0.65 and non-zero slope is significant at p<0.001 for within-session data plot).

### Starting Point identification for evaluating progression of differential T1/T2 refinement

To more precisely examine whether proximal action (T2) refinement precedes that of distal action (T1) in Slow Learners, it was important to consider refinement progression of T1 relative to T2. To rule out any bias towards proximal refinement because of initial bias towards proximal T2 refinement, a specific session was chosen as a Starting Point for analysis for each animal (Fig. 5c). This Starting Point is defined by an early session in which T1 and T2 were relatively similar in refinement levels or when the distal action T1 was more refined than proximal T2. To identify these Starting Points, a scan was made retrospective from the session for which the T1➔T2 time interval is close to final value (less than or equal to a median of 3 seconds). Using this approach, we identified earlier sessions in which distal T1 refinement was equal to or greater than proximal T2 (T2 – T1 refinement curve area less than or equal to 0). The latest such session was set as the Starting Point for analysis. If at no point early in learning did an animal have a session where proximal (T1) action is most refined relative to distal (T2) action, an early session of closest T1 and T2 refinement was used as the Starting Point. The initial T2-T1 refinement curve area difference calculated from the Starting Point to next session was subtracted from all T2-T1 area differences calculated in subsequent sessions. This value is called the Starting Point subtracted refinement difference (Fig. 5c). This made it possible to clearly track the change in relative refinement of distal(T1) vs. proximal(T2) actions over time (Values above zero indicate T2>T1 refinement, and values below zero indicate T1>T2 refinement). To identify the Turning Points for each animal, sessions carrying the local maximum value of the Starting Point subtracted refinement difference were identified for each animal. To calculate Starting Point subtracted refinement, scaled refinement values from sessions of interest were subtracted from that of the Starting Point session defined above (Fig. 5g).

### Odds ratio analysis

For odds ratio calculation (Fig. 5e), the total amount of open field ➔ Turning Point session (second of two consecutive sessions used to calculate the refinement difference at Turning Point as mentioned above) and Turning Point ➔ session of criterion frequency median interval changes was summed up for T1➔T2 and T2➔T1 intervals, respectively. Next, the proportion of total interval change stemming from the open field condition➔Turning Point period, and from Turning Point➔session reaching criterion frequency period, were calculated. Next, the proportion of open field➔Turning Point interval change was divided by the proportion of Turning Point ➔ session reaching criterion frequency period interval change for T1➔T2 and T2➔T1 interval types, respectively. This gives the odds ratio.

### T1 probability rank change across time bins from T2 trigger

For every actual or simulated trigger for T1➔T2 performance, the first occurrences of every action before or after T2 triggers were counted at specific 300 ms time bins for up to 6 seconds before and after T2 trigger (Fig. 5h). This was done for the specific conditions of baseline, Starting Point, Turning Point, session passing criterion frequency, and last session. The probability of an action occurring at a specific 300 ms time bin was calculated for all actions in the repertoire, and the values were used to determine probability rank in terms of percentiles (100 percentile is most probable action relative to all actions at a specific 300 ms time bin). To assess total T1 probability rank change within 0.3-1.8 or 2.1-3.6 second time bins, the area under the curve was determined and values were normalized by subtraction from each animal’s corresponding baseline values.

### Statistical Analysis

Standard statistical analyses were performed on Prism (Versions 7, 9, 10; GraphPad Software, Inc.; RRID:SCR_002798) and permutation/bootstrap analyses were performed on MATLAB (MathWorks Inc, RRID:SCR_001622). To determine appropriate tests for comparisons, datasets were assessed for normality using Anderson-Darling, D’Agostino & Pearson, Shapiro-Wilk and/or Kolmogorov-Smirnov tests whenever applicable. Datasets were also visualized for normality using QQ plots and assessed for equal variance by examining the Residual plot (Residuals vs. Predicted Y). Parametric or non-parametric tests were chosen based on the combination of these analyses. Data were transformed logarithmically (with or without addition of a constant prior to transformation) whenever it was appropriate to promote normality and equal variance. Unless specified, sphericity was not assumed, and Geisser-Greenhouse correction was applied in all ANOVA tests. The appropriate post hoc multiple comparisons tests were applied to compare between the means of specific conditions wherever applicable. Significance was set at alpha = 0.05. For bootstrap analysis, significance was determined by asking whether the original target action mean Fano factor was greater or less than the 95% confidence interval of the bootstrap distribution. Permutation test was applied in the comparisons between regression models because of the large sample size discrepancy between groups. Bonferroni p adjustment was used to account for multiple comparisons in this case. When Graphpad Prism does not output exact p-value, Excel (version 16.78.3; Microsoft) was used with the ANOVA-specific FDIST(F, DFN, DFD) where DFN is the Numerator Degrees of Freedom and DFD is the Denominator Degrees of Freedom. For detailed description of statistical procedures please refer to Supplementary Information.

## Acknowledgements

We thank V.Athalye for helpful feedback, A. Vaz and C. Carvalho for mouse colony management, members of the Costa laboratory for comments, and the help from the Scientific Hardware Platform, Histopathology Platform, Scientific Software Platform and Advanced Bioimaging & BioOptics Experimental Platform (member of the Portuguese Platform of Bioimaging (PPBI-POCI-01-0145-FEDER-022122) of the Champalimaud Institute, S. Mutlu, D. Bento, P. Carriço., P. Silva, J. Araujo for hardware/software assistance, I. Marcelo for assistance with multinomial logistic regression, N. Loureiro for assistance with inertial sensor calculations, M. Mendoça, C. Alcacer, H. Rodrigues and D. Peterka for help with experimental setups. M. Carey lab for sharing apparatus. S. Fusi for project feedback. R. Junker for advice on dynamic range boxes. This work was supported by Life Sciences Research Fellowship and NINDS K99/R00 Award (1K99NS112575) granted to J.C.Y.T and National Institute of Health funding (5U19NS104649) to R.M.C.

## Author Contributions

We thank V.Athalye for helpful feedback, A. Vaz and C. Carvalho for mouse colony management, members of the Costa laboratory for comments, and the help from the Scientific Hardware Platform, Histopathology Platform, Scientific Software Platform and Advanced Bioimaging & BioOptics Experimental Platform (member of the Portuguese Platform of Bioimaging (PPBI-POCI-01-0145-FEDER-022122) of the Champalimaud Institute, S. Mutlu, D. Bento, P. Carriço., P. Silva, J. Araujo for hardware/software assistance, I. Marcelo for assistance with multinomial logistic regression, N. Loureiro for assistance with inertial sensor calculations, M. Mendoça, C. Alcacer, H. Rodrigues and D. Peterka for help with experimental setups. M. Carey lab for sharing apparatus. S. Fusi for project feedback. R. Junker for advice on dynamic range boxes. This work was supported by Life Sciences Research Fellowship and NINDS K99/R00 Award (1K99NS112575) granted to J.C.Y.T and National Institute of Health funding (5U19NS104649) and the Aligning Science Across Parkinson’s (ASAP-020551) through the Michael J. Fox Foundation for Parkinson’s Research (MJFF) to R.M.C. The authors have applied a CC BY NC public copyright license for the manuscript.

## Additional Information

Supplementary Information is available for this paper.

## Code availability

MATLAB (MathWorks) codes used for data analysis are available from the corresponding author.

## Data availability

Source Data are available from the corresponding author are deposited onto Zenodo (DOI: 10.5281/zenodo.10146089) upon reasonable request.

Correspondence and requests for materials should be addressed to

## Extended Data Figure Legends

**Extended Data Figure 1.**
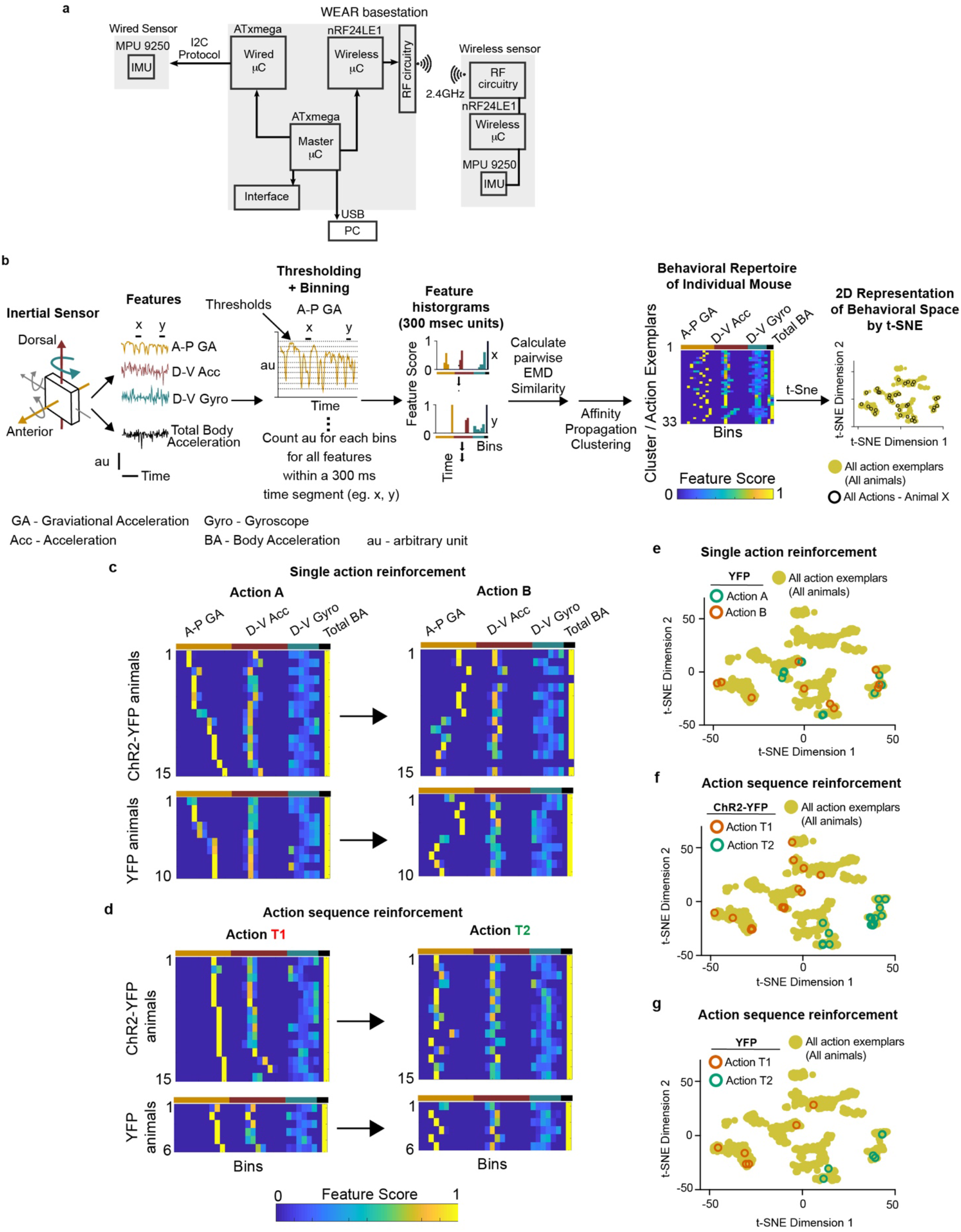
Behavioral monitoring hardware, action clustering and selected actions for reinforcement. **a,** Schematic showing the WEAR basestation and communications with either wired (not used in this study) or wireless inertial sensors. **b,** Detailed schematic showing processing of inertial sensor data for behavioral clustering and 2-D behavioral repertoire visualization. **c,** Heat map showing motion feature score distributions of exemplar Action A (left) and Action B (right) reinforced for each animal. **d,** Heat map showing motion feature score distributions of exemplar Action T1 (left) and Action T2 (right) reinforced for each animal. **e-g,** t-SNE plots showing 2-D behavioral space showing all action exemplars pooled from all animals (yellow circles) overlaid with actions targeted in single action reinforcement YFP (e), two action ChR2-YFP (f) and YFP (g) animals.

**Extended Data Figure 2.**
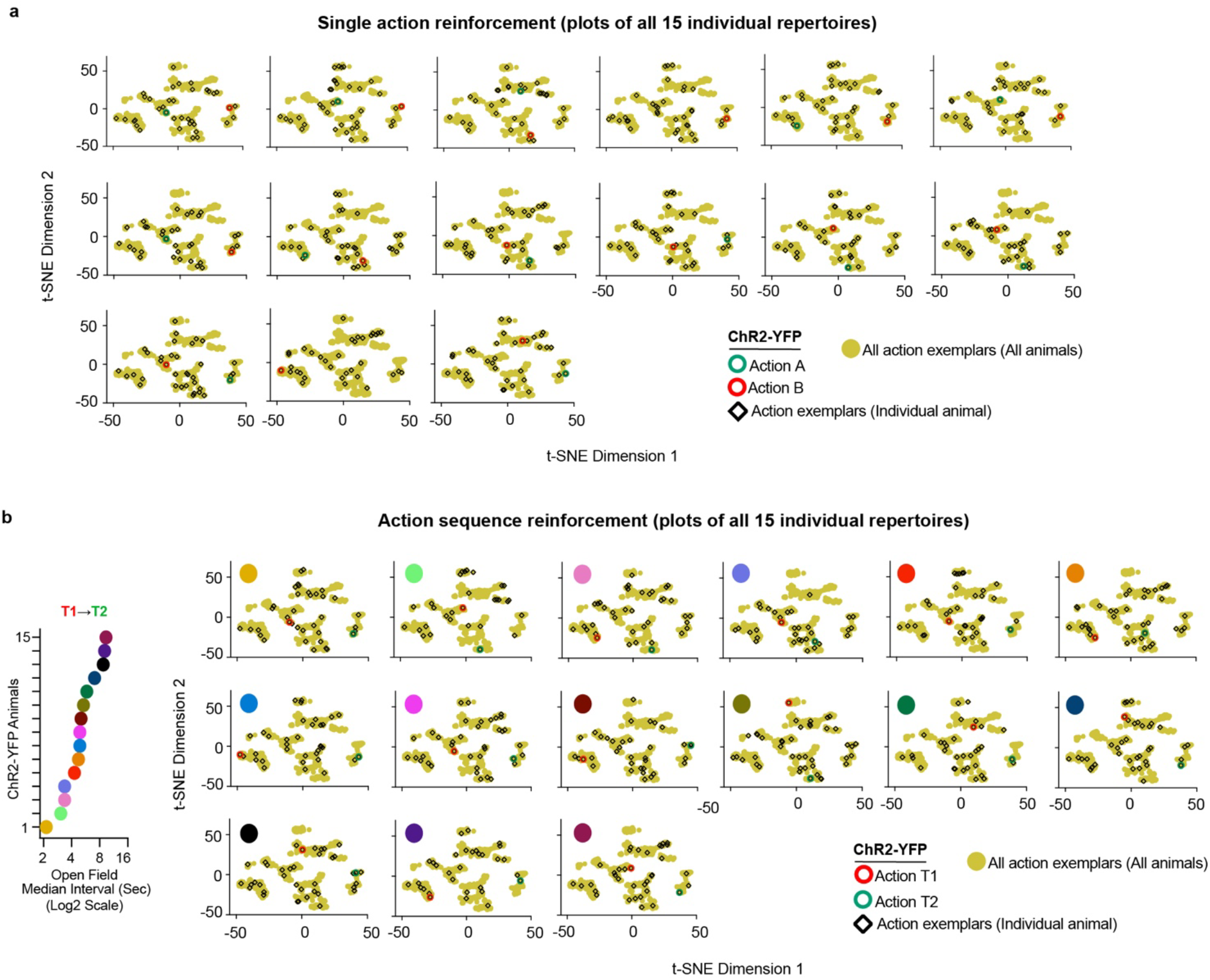
Behavioral spaces of individual ChR2-YFP animals used in this study. **a-b,** t-SNE plots showing distribution of all action exemplars in individual ChR2-YFP animals across the 2-D behavioral space (yellow circles, pooled from all animals) in single action (**a**) and action sequence (**b**) learning. **b,** The plots of action sequence learners are ordered according to their initial open field median T1→T2 intervals. Leftmost graph: Open field inter-action intervals of T1/T2 pairs chosen. Same color codes in right t-SNE graphs. n = 15 ChR2-YFP animals (biological replicates) in each experiment.

**Extended Data Figure 3.**
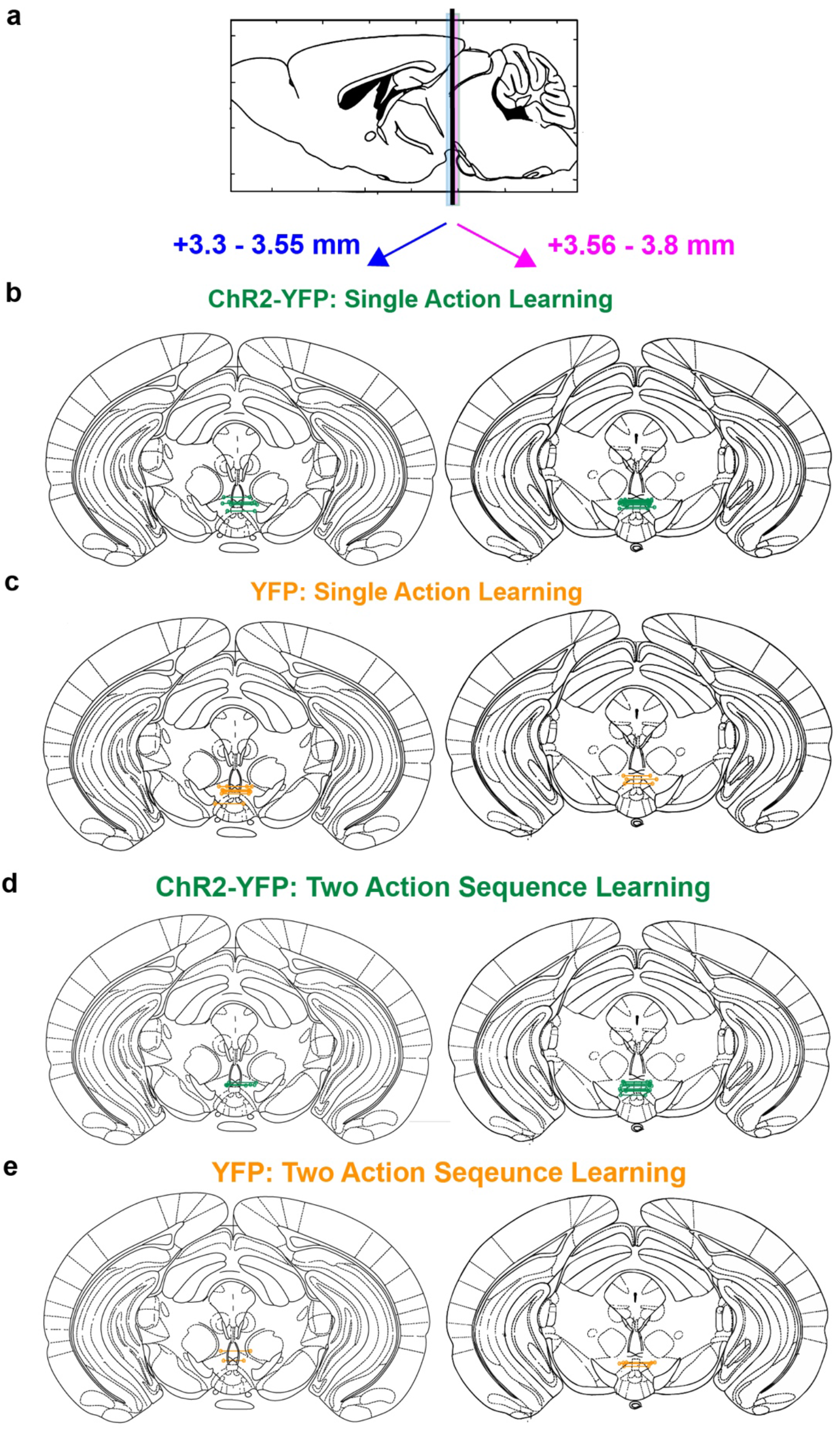
Implant locations for all animals used in this study. **a,** Sagittal view of mouse brain with labeling of anterior (blue) and posterior (magenta) brain regions surrounding injection/implant coordinates (vertical black line). **b-c,** Actual dual cannula implant locations of individual ChR2-YFP (*n*=15; green) (**b**) and YFP (*n* = 10; orange) (**c**) animals used for single action reinforcement experiments. (n: animals - biological replicates) **d-e,** Actual dual cannula implant locations of individual ChR2-YFP (*n*=15; green) (**d**) and YFP (*n* = 6; orange) (**e**) animals used for two action sequence reinforcement experiments. Individual dual implant locations are marked by open circles joined by horizontal lines.

**Extended Data Figure 4.**
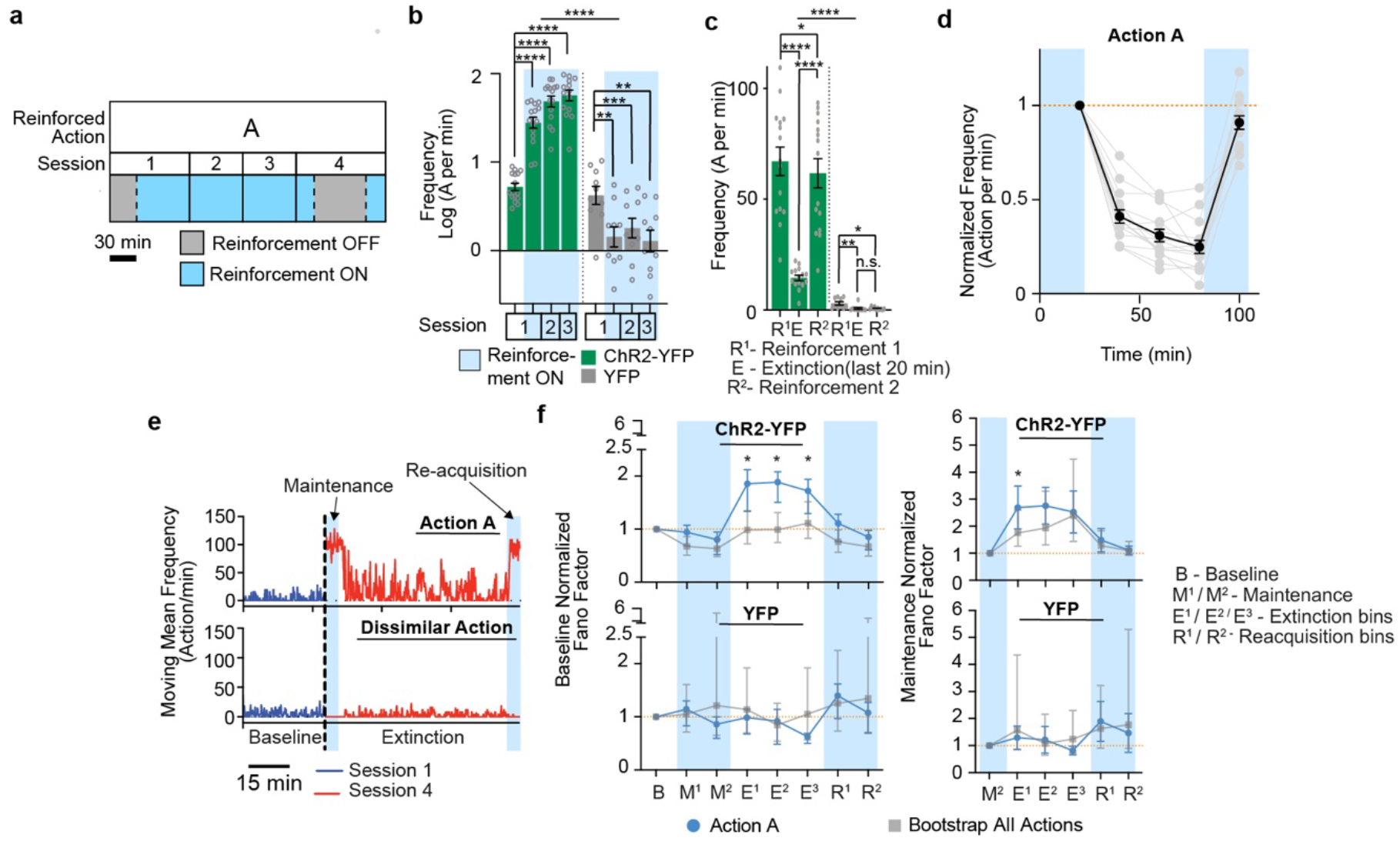
Reinforcement of action A in ChR2-YFP and YFP animals. **a-d,** *n* = 15 ChR2-YFP, 10 YFP animals (biologically independent). Plots in (**b-d**) show mean/S.E.M. **a,** Protocol for Action A reinforcement. **b-c.** Repeated measures two-way ANOVA reveal significant difference across time and ChR2-YFP/YFP groups in reinforcement sessions (F(3,69) = 82.61, p < 0.0001) and extinction session (F(2,46) = 18.66, p < 0.0001). Tukey’s post hoc comparisons. **d,** Detailed extinction of action A reinforcement showing mean (black) and individual curves (light grey lines). Plot shows means/S.E.M. Asterisks: **** p < 0.0001. *** p < 0.001. ** p < 0.01. * p < 0.05. n.s. – not significant. **e**, Bursty target action behavior upon extinction. Plots show moving mean frequencies of action A (target action) and a dissimilar action of an ChR2-YFP animal over baseline and extinction conditions. The dissimilar action is later reinforced for and is referred to as action B in subsequent protocols. **f,** Normalized Fano factor over extinction conditions (action A). Plots were mean/95% confidence intervals. *n* = 15 ChR2-YFP, 9 YFP animals. * - Mean of ChR2-YFP animals outside bootstrapped 95% confidence intervals; 2-tailed comparison. See Supplementary Information for statistical/sample details.

**Extended Data Figure 5.**
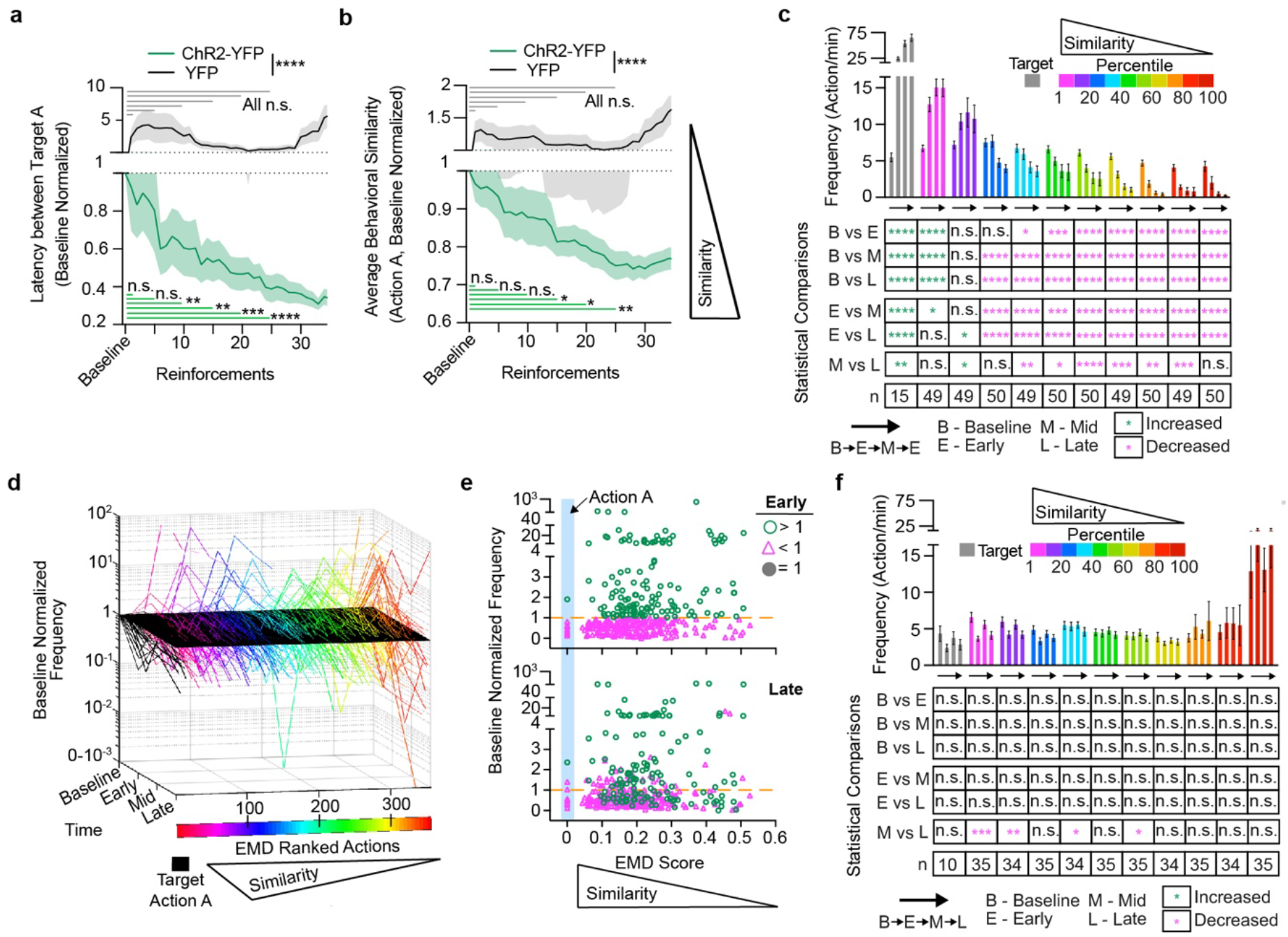
Reinforcement of Action A in ChR2-YFP and YFP animals. **a-f,** *n* = 15 ChR2-YFP, 10 YFP animals (biological replicates). Plots in (**a-c, f**) show mean/S.E.M. **a-b,** Rapid behavioral adjustments to close-loop reinforcements. Repeated-Measures 2way ANOVA, posthoc 2-sided Sidak’s multiple comparisons test: Significant Time x Group Interactions (F(35, 805) = 3.019, p<.0001 (**a**); F(35, 805)=3.086, p<0.0001 (**b**)). **c,** Expanded graph from Figure 1 showing quantification of raw frequency changes across learning stage and target similarity percentile groups. Additional table below graph indicate 2-tailed, Tukey multiple comparisons results. *n* – sample size (actions). **d,** Evolution of pooled action repertoire (n = 355 actions from 10 YFP animals) across action learning from a naïve state. A black parallelogram was set at baseline normalized frequency = 1 to help visualization. **e,** Cross-sectional view of action repertoire frequencies at Early and Late stages. **f,** Same as (**c**) but for YFP animals. Asterisks: **** p < 0.0001. *** p < 0.001. ** p < 0.01. * p < 0.05. n.s. – not significant. See Supplementary Information for statistical/sample details.

**Extended Data Figure 6.**
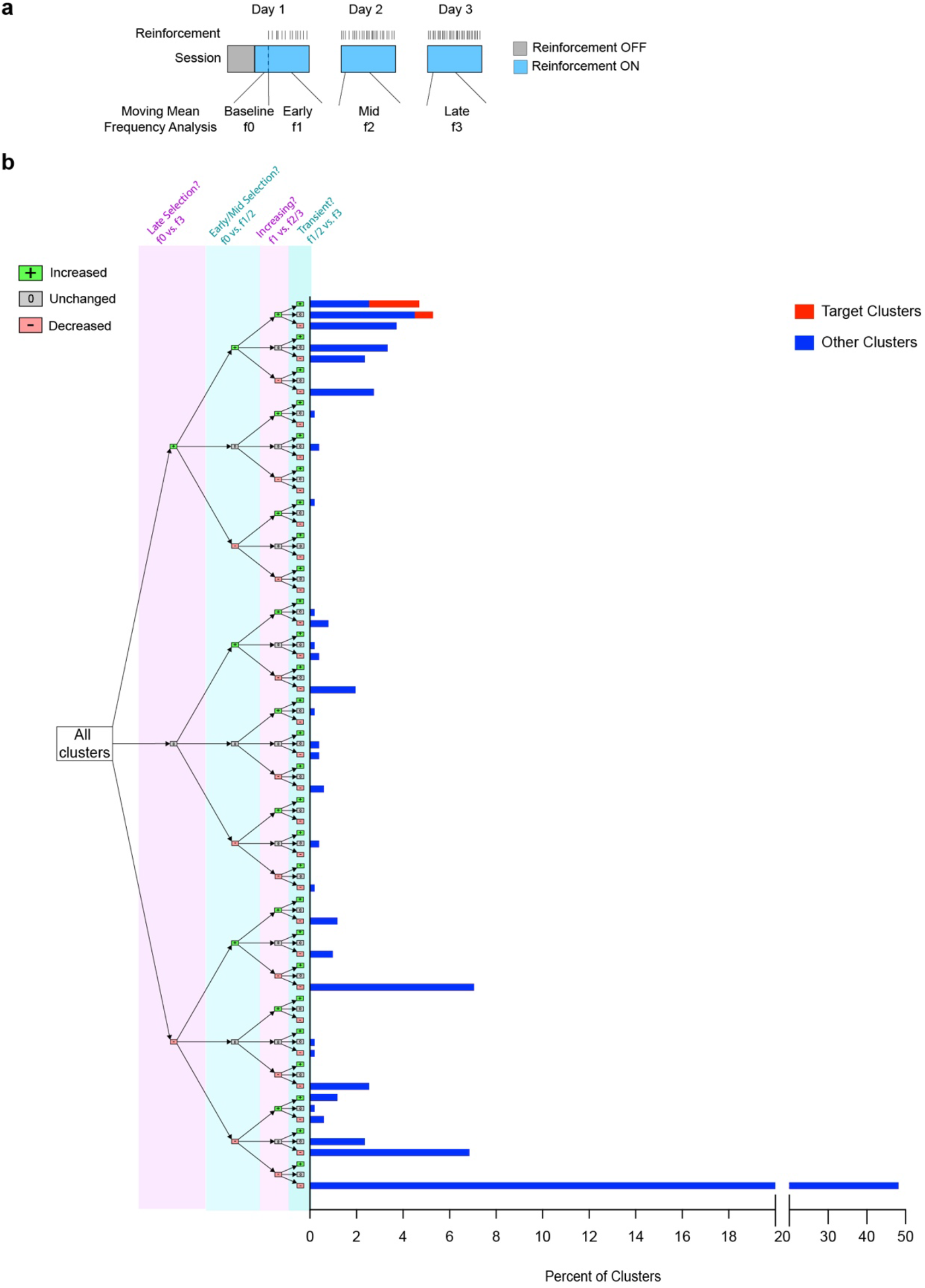
Classification of action dynamic types upon Action A reinforcement. **a,** Schematic of reinforcement protocol and definition of learning stages. f0, f1, f2 and f3 represent the moving mean frequencies from Baseline, Early, Mid and Late stages, respectively. **b,** Distribution of all actions according to criterion. 4 criteria were used. First, actions were divided by their final modulation status (Late Selection? f0 vs. f3). The next division is based on whether they were significantly modulated in early stages (Early/Mid Selection? f0 vs. f1/2). The third division is based on comparing initial modulation status in Early stage with later stages (Increasing? f1 vs. f2/3). The final division is based on whether actions in early stages decrease significantly by the Late stage (Transient? f1/2 vs. f3). Distribution across all possible combination of these comparison outcomes is plotted in terms of Percent of All Actions on the x-axis. *n* = 511 actions (496 other actions (blue bars), 15 target Actions A (red bars)), 15 ChR2-YFP animals (biological replicates).

**Extended Data Figure 7.**
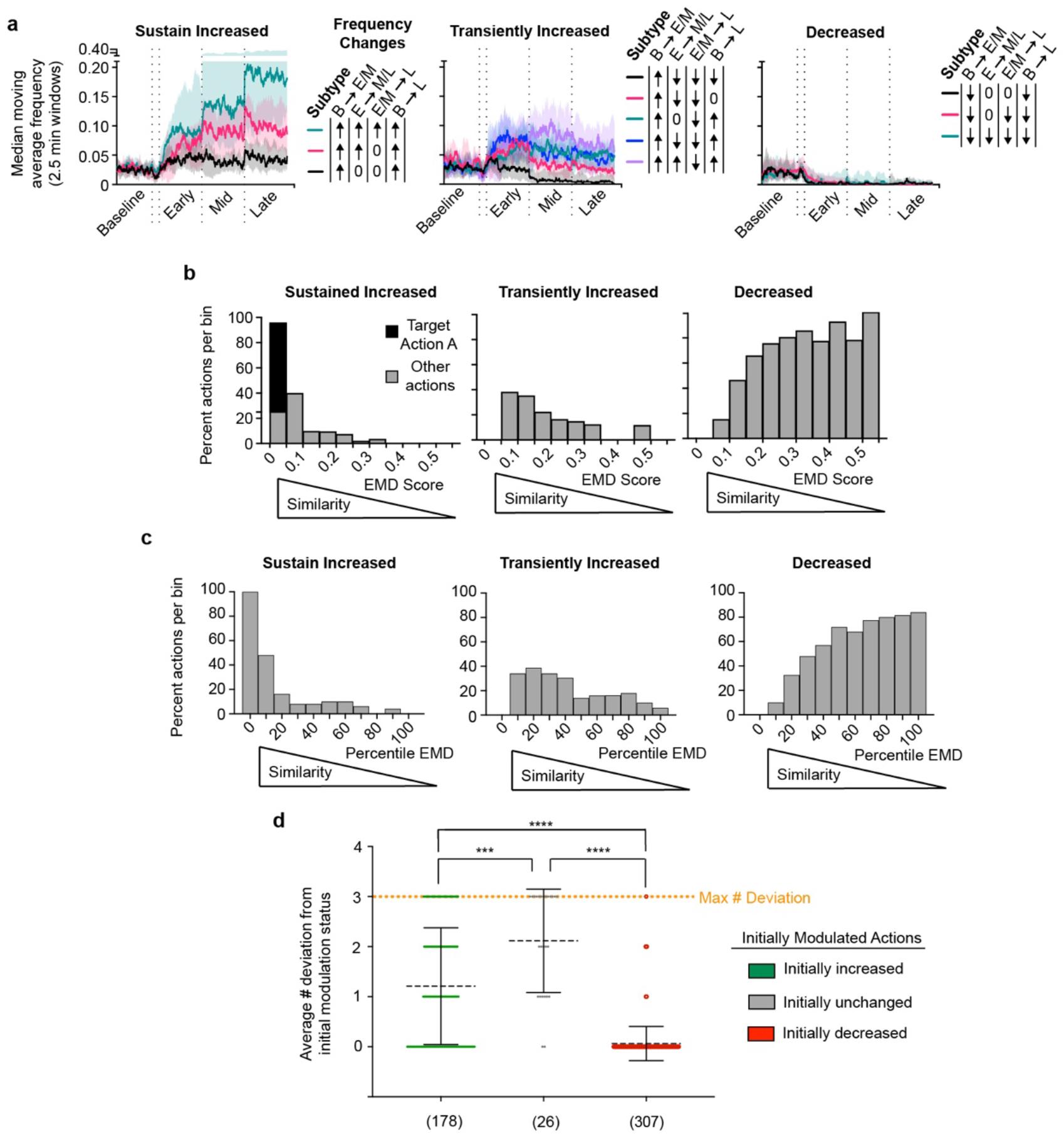
Classified Action Dynamic types and analyses. **a-d**, n = 15 ChR2-YFP, 10 YFP animals (biological replicates). Plots in (d) show mean/S.E.M. **a,** Three defined types of action dynamics. Plots were median moving average frequency of subtypes (lines) and 25th-75th percentile ranges (color filled). Significant modulation: +, increased. -, decreased. 0, unchanged. **b**, Distribution of action dynamic types shift according to their percentile similarity to target. Histograms show percent clusters per bin vs. target action similarity. **c,** Distribution of action dynamics types shift according to action similarity to target. **d,** Initially decreased actions tend to retain its decreasing or decreased status throughout learning. Plot shows mean/s.t.d. (Kruskal-Wallis test, medians vary significantly, p < 0.0001. Kruskal-Wallis Statistic 237.8. 2-tailed Dunn’s post hoc tests: Initially increased (n = 178 actions) vs. decreased (n = 307 actions) – p < 0.0001. Initially increased vs. unchanged (n = 26 actions) – p = 0.0004. Initially decreased vs. unchanged – p < 0.0001). Asterisks: **** p < 0.0001. *** p < 0.001. ** p < 0.01. * p < 0.05. n.s. not significant. See Supplementary Information for details.

**Extended Data Figure 8.**
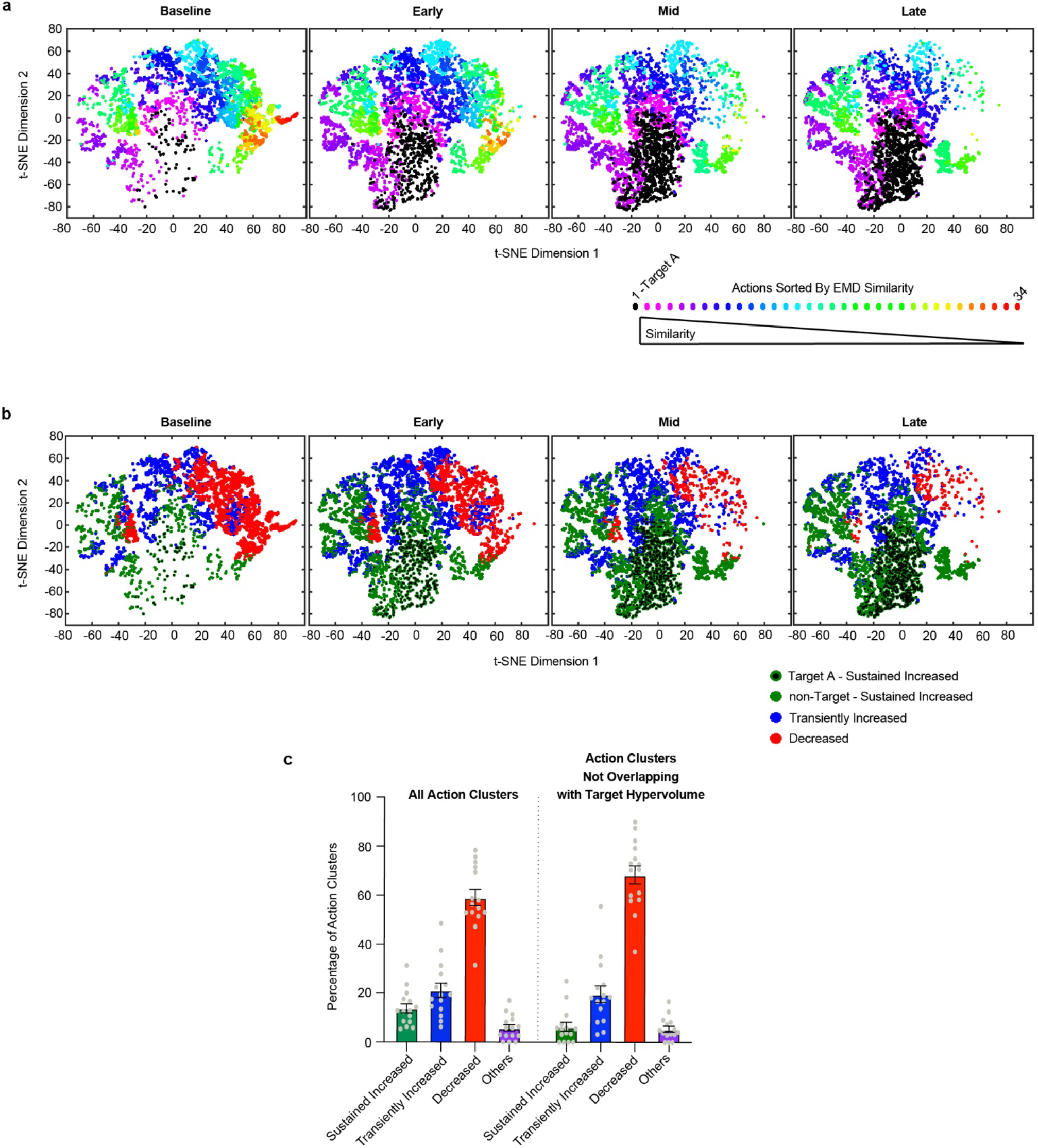
t-SNE and hypervolume analysis supplementing Figure 1. **a-b,** t-SNE representing the evolution of actions over learning to perform Target A. Shown is the behavioral evolution of a representative ChR2-YFP animal. Actions clusters are color labeled according to Earth Mover’s Distance (EMD) similarity to target action in (**a**) and according to action dynamics type in (**b**) (34 action clusters in total). Dots represent single action instances. Black dots (**a**) and black dots with green outlines (**b**) represent single instances of Target A. **c,** Percentage of classified action dynamic types based on all action clusters (left set of bars) or action clusters whose hypervolume did not overlap with target hypervolume (right set of bars). Bar graphs indicate mean +/- S.E.M. n = 15 ChR2-YFP animals (biological replicates).

**Extended Data Figure 9.**
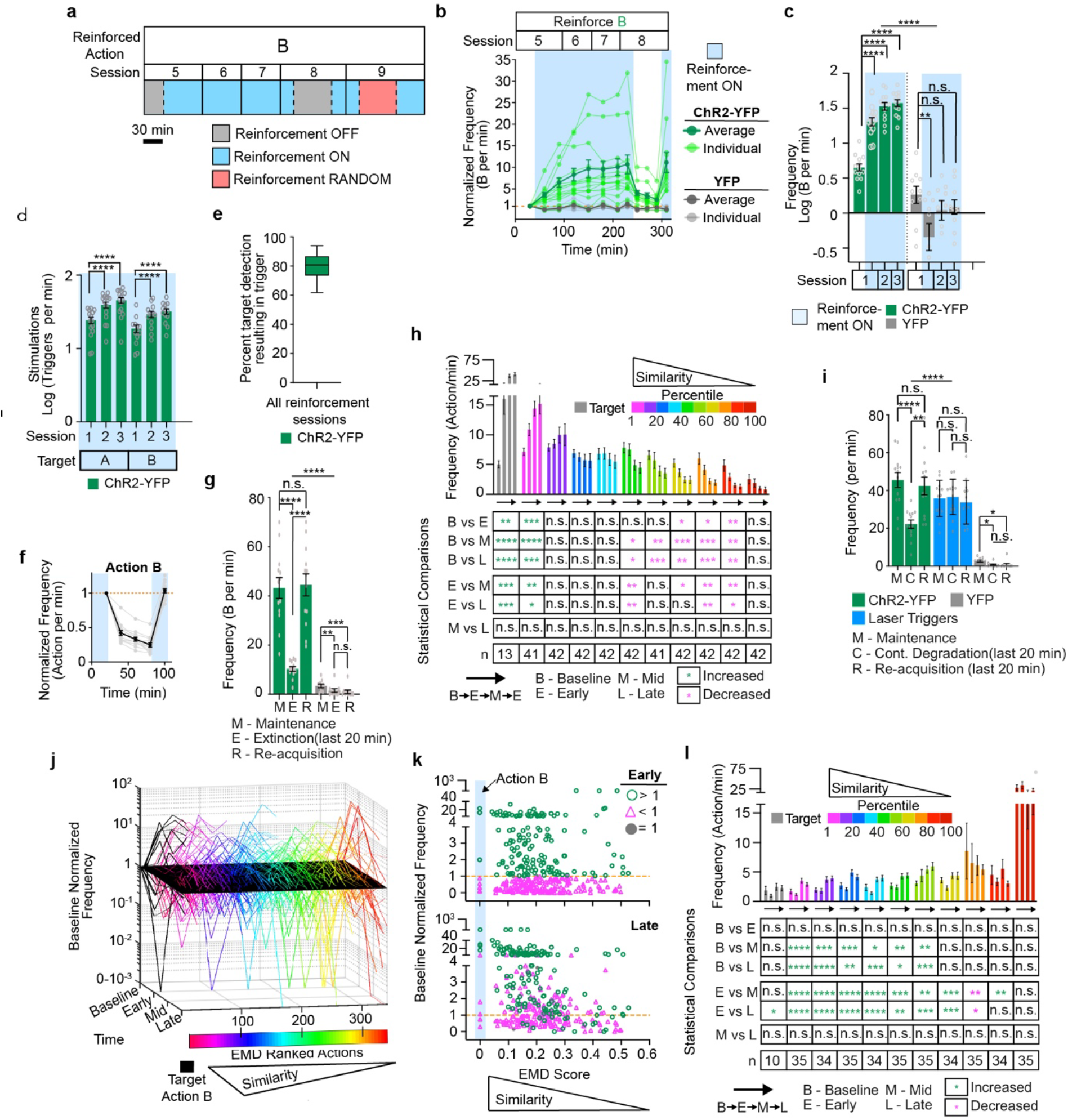
Closed-loop reinforcement of Action B in ChR2-YFP and YFP animals. **a-l**, n = 13 ChR2-YFP (Action B), 15 ChR2-YFP (Action A), 10 YFP (Action B) animals (biological replicates). All bar and line plots (**b-d, f-i, l**) show mean/S.E.M. **a,** Protocol for Action B reinforcement, extinction and contingency degradation. **b,** ChR2-dependent increase in Action B. **c,** Quantification of (**b**). ChR2-YFP and YFP groups differed across time and groups (Two-way ANOVA. F(3,63) = 38.67, p<0.0001). Tukey’s post hoc comparisons. **d,** Actual stimulation frequency for single action learning. Significant increases across time per group (1-way ANOVA. Action A: F(1.34, 18.8)=112, p<0.0001. Action B: F(1.18, 14.2) = 46.5, p<0.0001). Two-sided Dunnett multiple comparisons test. **e,** Percent target detection resulting in trigger. Tukey plot (median/25^th^-75^th^ percentile box/min-max fences). **f,** Extinction and re-acquisition of Action B showing mean (black) individual curves (light grey lines). **g,** Quantification of (**f**). ChR2-YFP and YFP groups differed across time and groups (Two-way ANOVA. F(2,42) = 32.23, p < 0.0001). ChR2-YFP and YFP animals decreased Action B frequency by the end of extinction, but only ChR2-YFP animals significantly increased Action B frequency by the end of re-acquisition period. 2-sided Tukey’s post hoc multiple comparisons test. **h,** Quantification of raw frequency changes across learning stage and target similarity percentile groups in ChR2-YFP animals. The p value results of two-way mixed effect model analysis is shown in the table. **i**, Quantification of contingency degradation results. Actin B and Laser Trigger frequencies differ across time and groups (Two-way ANOVA. F(2,48) = 26.19, p < 0.0001). 2-sided Tukey’s post hoc multiple comparisons test. **j**, Evolution of pooled action repertoire (n = 341 actions from 10 YFP animals) across action learning from a naïve state. A black parallelogram was set at baseline normalized frequency = 1 to help visualization. **k,** Cross-sectional view of action repertoire frequencies at Early and Late stages. **l,** Same as (**h**) but for YFP animals. See Supplementary Information for statistical/sample details.

**Extended Data Figure 10.**
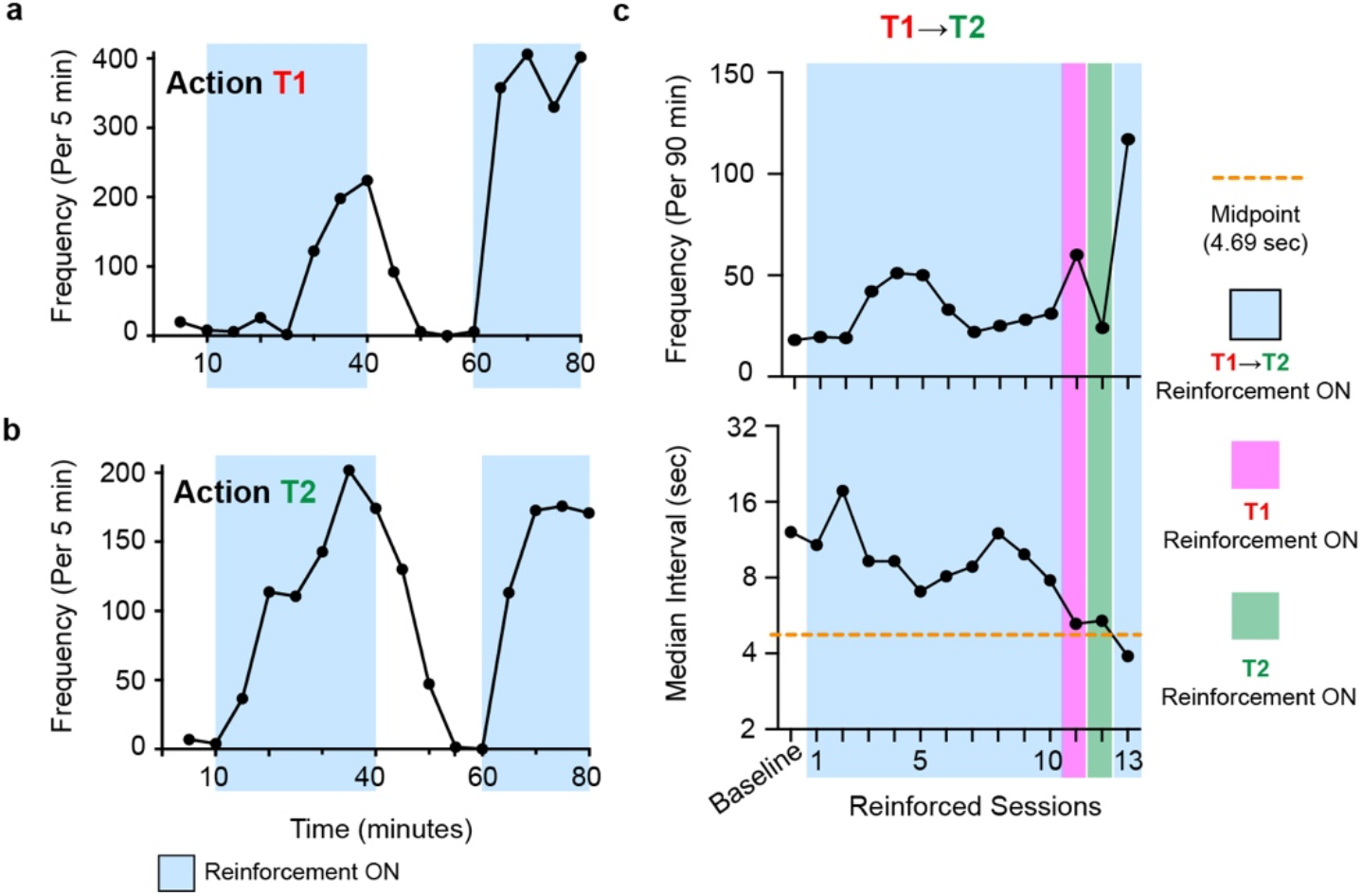
Two-action learning analyses supplementing Figure 4. **a-c,** Individual ChR2-YFP animal that failed to reach criterion frequency by session 10 and had the largest T1➔T2 median interval of all animals at session 10 was tested for the ability to be reinforced for single actions T1 (**a**) and T2 (**b**) in order. Upon verifying that the individual could learn T1 and T2 separately in a ChR2-YFP manner (**a, b**), the animal was subjected again to T1➔T2 reinforcement (**c**). **c,** Increased T1➔T2 performance (top) coincided with decreased T1➔T2 median interval (bottom) below the sigmoidal midpoint (orange dashed lines) fitted in Fig. 4l.

**Extended Data Figure 11.**
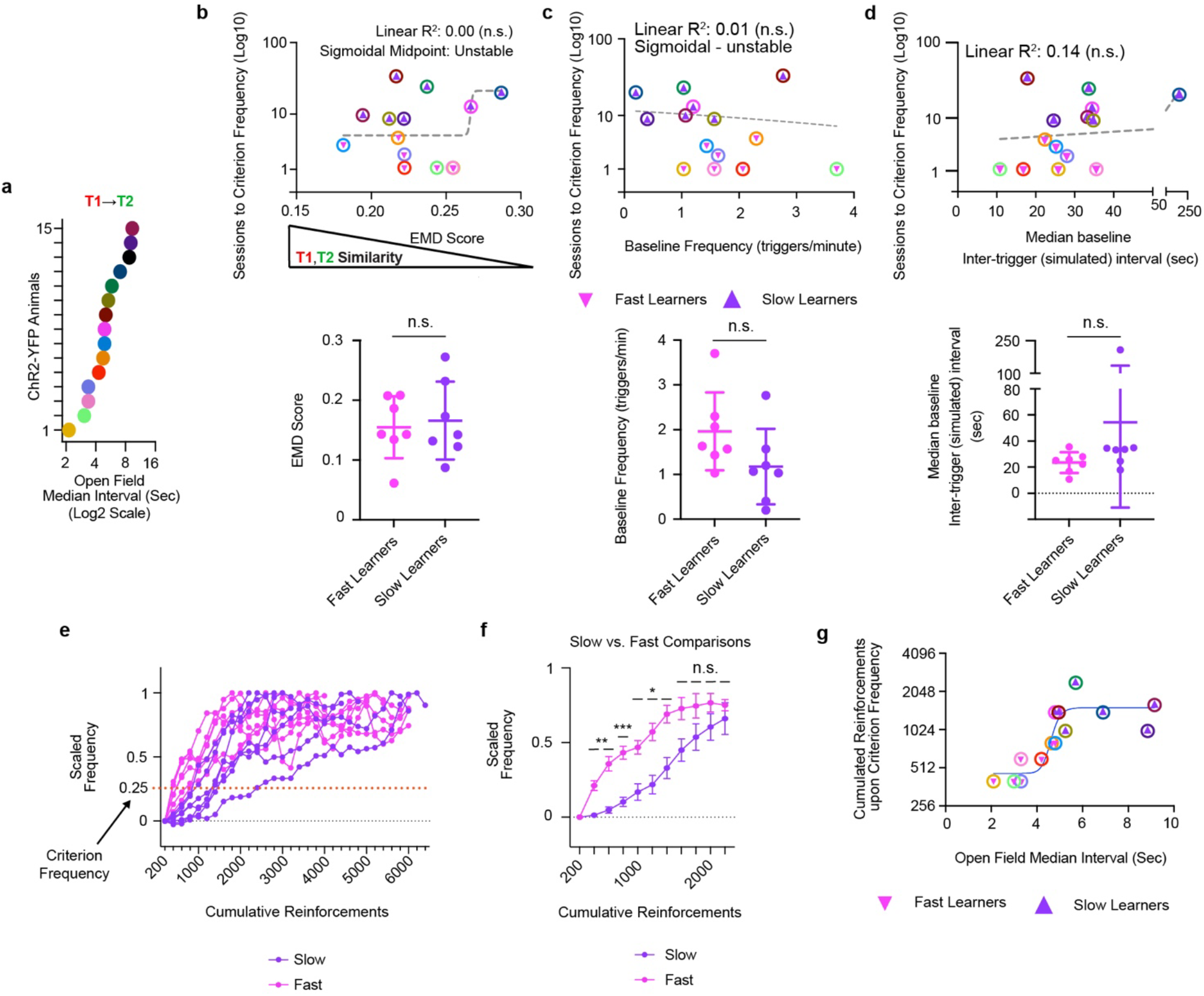
Two-action learning analyses supplementing Figure 4. **a,** Open field inter-action intervals of T1/T2 pairs chosen. Same color codes in (**b-d, f**). **b-d,** Top panels: Action similarities between T1 and T2 (**b**) baseline T1 ➔ T2 frequencies (simulated triggers per minute) (**c**) and median baseline inter-simulated trigger intervals (**d**) do not account for variation in sessions to criterion frequency. (F-tests: F(1,12) = 0.04(**b**), 1.35(**c**), 1.91(**d**): p=0.85(**b**), p=0.27(**c**), p=0.19(**d**). Grey dashed line – model fit (sigmoidal in (**b**) and linear in (**c,d**). Bottom panels: No significant difference between groups in any independent variables shown in the top panels (p=0.73 (**b**), p=0.11 (**c**), p = 0.24 (**d**)). Mean +/- s.t.d. 2-tailed, unpaired t-test assuming equal variance. **e,** Individual learning curves scaled to initial session frequency (value of 0) and maximum frequency (value of 1). **f,** Slow learners still show significantly slower learning after matching for number of reinforcements. Plot shows Mean/S.E.M. Significant interaction between cumulative reinforcement bins and groups (2-way ANOVA. F(10, 120) = 5.35) p<0.0001). 2-tailed Sidak’s multiple comparisons (left to right comparisons on graph: p= 0.0004, 0.002, 0.0005, 0.016, 0.016, 0.03, 0.24, 0.72, 0.86,1). **g**, Sigmoidal relationship between open field T1➔T2 interval and cumulated reinforcements upon criterion scaled frequency. n = 14 ChR2-YFP mice (biological replicates) (**b-g**). Asterisks: *** p < 0.001. ** p < 0.01. * p < 0.05. n.s. not significant. See Supplementary Information for statistical/sample details.

**Extended Data Figure 12.**
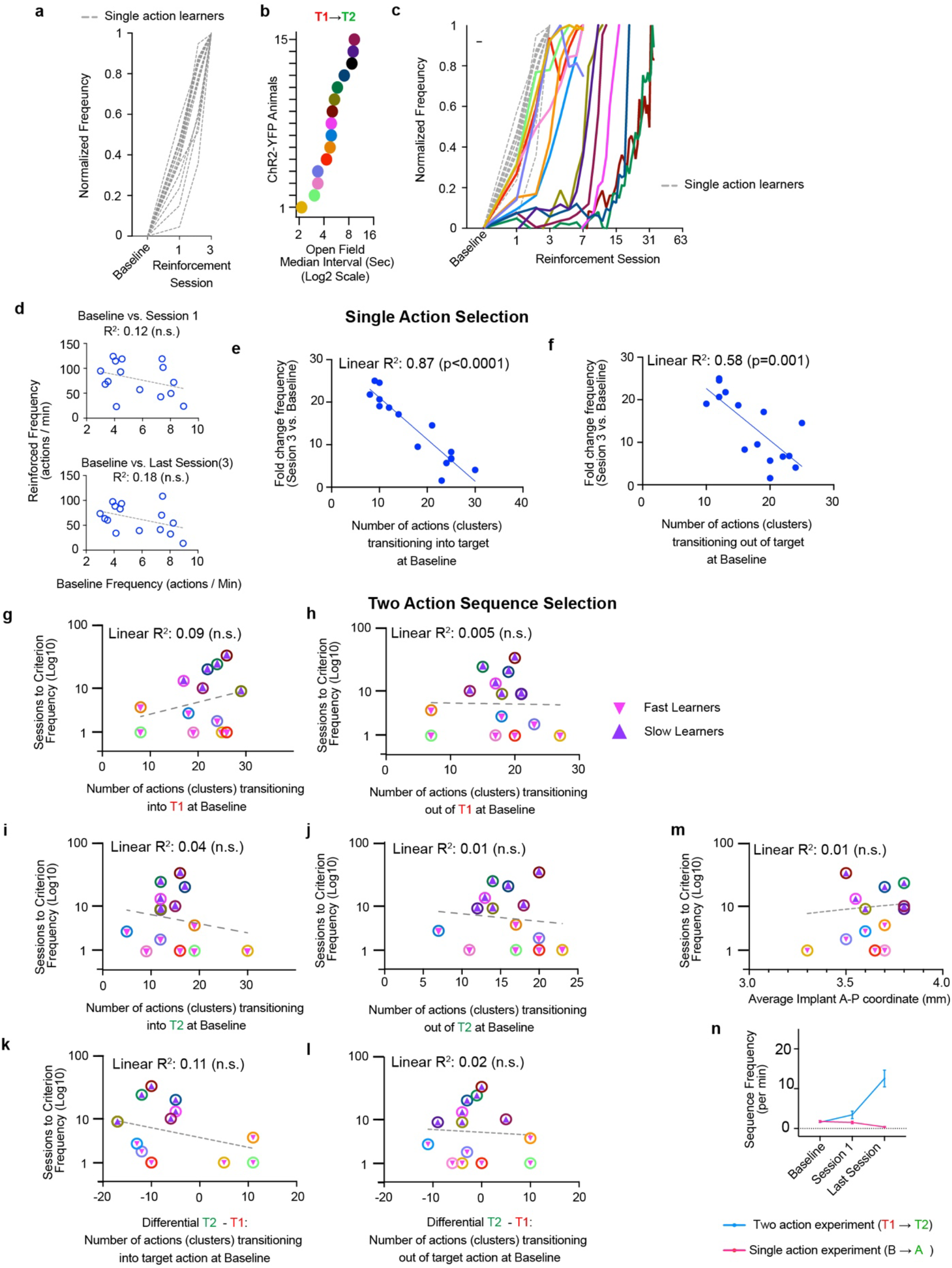
Two-action learning analyses supplementing Figure 4. **a-c,** Emergence of distinct type of learning curves specifically amongst action sequence learners. **a,** Learning curves of single action learners. Normalized frequency indicates that frequencies were normalized to the highest frequency achieved in a session for each ChR2-YFP animal. **b,** Open field inter-action intervals of T1/T2 pairs chosen in action sequence learning. Same color codes in (**c, g-m**). c, Overlaying learning curves of single action learners (grey, dashed lines) with action sequence learners (color coded as in (**b**)). **d-f,** Linear relationships assessed in single action learners. **d,** No significant linear relationship between baseline action frequency and reinforced frequencies in Session 1 (top) and 3 (bottom). **e-f,** The number of action (clusters) transitioning into (**e**) and out of (**f**) target at baseline is negatively related to the fold change in frequency from baseline to last session. F-test (F(1,13)=83.31(**e**), 17.88(**f**); p=5*10^-7(**e**),1.0*10^-3(**f**). **g-l,** Action-to-Target (**g, i**) and Target-to-Action (**h,j**) and differential T2-T1 (**k,l**) transitioning parameters are not related to sessions to criterion frequency parameter. F-test: F(1,12)=1.11(**g**), 0.0053(**h**), 0.037(**i**), 0.014(**j**),1.49(**k**),0.048(**l**); p=0.31(**g**), 0.94(**h**), 0.51(**i**), 0.69(**j**), 0.24(**k**), 0.83(**l**). Some data points overlap. **m,** No significant linear relationship between implant location and sessions to criterion. F-test: F(1,12)=0.11, p=0.75). Some data points overlap (**g-m**). **n,** Lack of sequence reinforcement in single action reinforcement. Learning curves differ across time and group (2-way ANOVA – F(2,54) = 143.3, p<0.0001). Plot indicates mean +/- SEM. n = 15 and 14 ChR2-YFP animals (biological replicates) in single action reinforcement and action sequence reinforcement (past criterion frequency) experiments, respectively. n.s. – not significant.

**Extended Data Figure 13.**
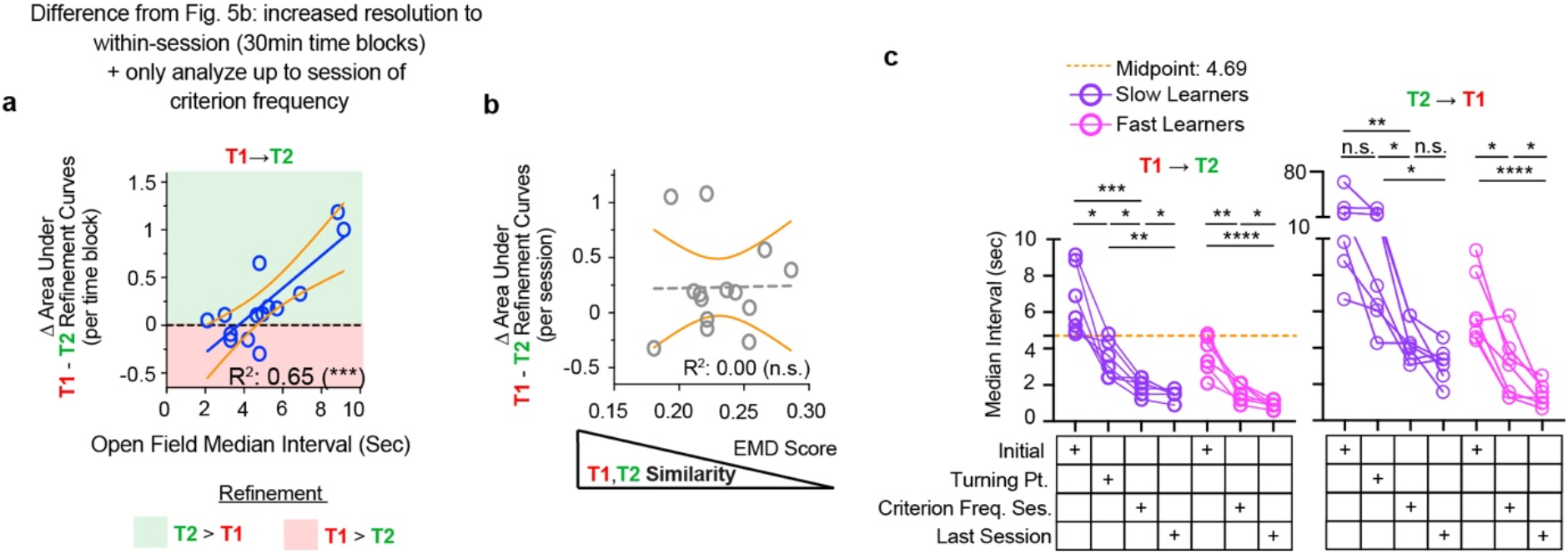
Two-action learning analyses supplementing Figure 5. **a,** Linear relationship between initial T1→T2 interval and differential T1-T2 refinement, despite accounting for within-session changes and excluding data upon reaching rising phase of learning. Non-zero slope significance: p < 0.001. **b,** Action similarities between T1 and T2 do not account for variation in differential T1-T2 refinement. n = 14 ChR2-YFP mice (biological replicates). Black dashed line - linear fit. Grey dashed line - sigmoidal fit. **c,** Full plots of T1➔T2 and T2➔T1 median interval changes across learning stages. Repeated-measures 2-way ANOVA. time-specific difference (Slow Learners – F(2.184, 26.20)=54.21, p=5.3*10^-10;Fast Learners – F(1.700, 20.40)=92.12, p=6.3*10^-9). Post-hoc 2-tailed Tukey’s multiple comparisons tests. **** p < 0.0001. *** p < 0.001. ** p < 0.01. * p < 0.05. n.s. – not significant. See Supplementary Information for statistical/sample details.

**Extended Data Figure 14.**
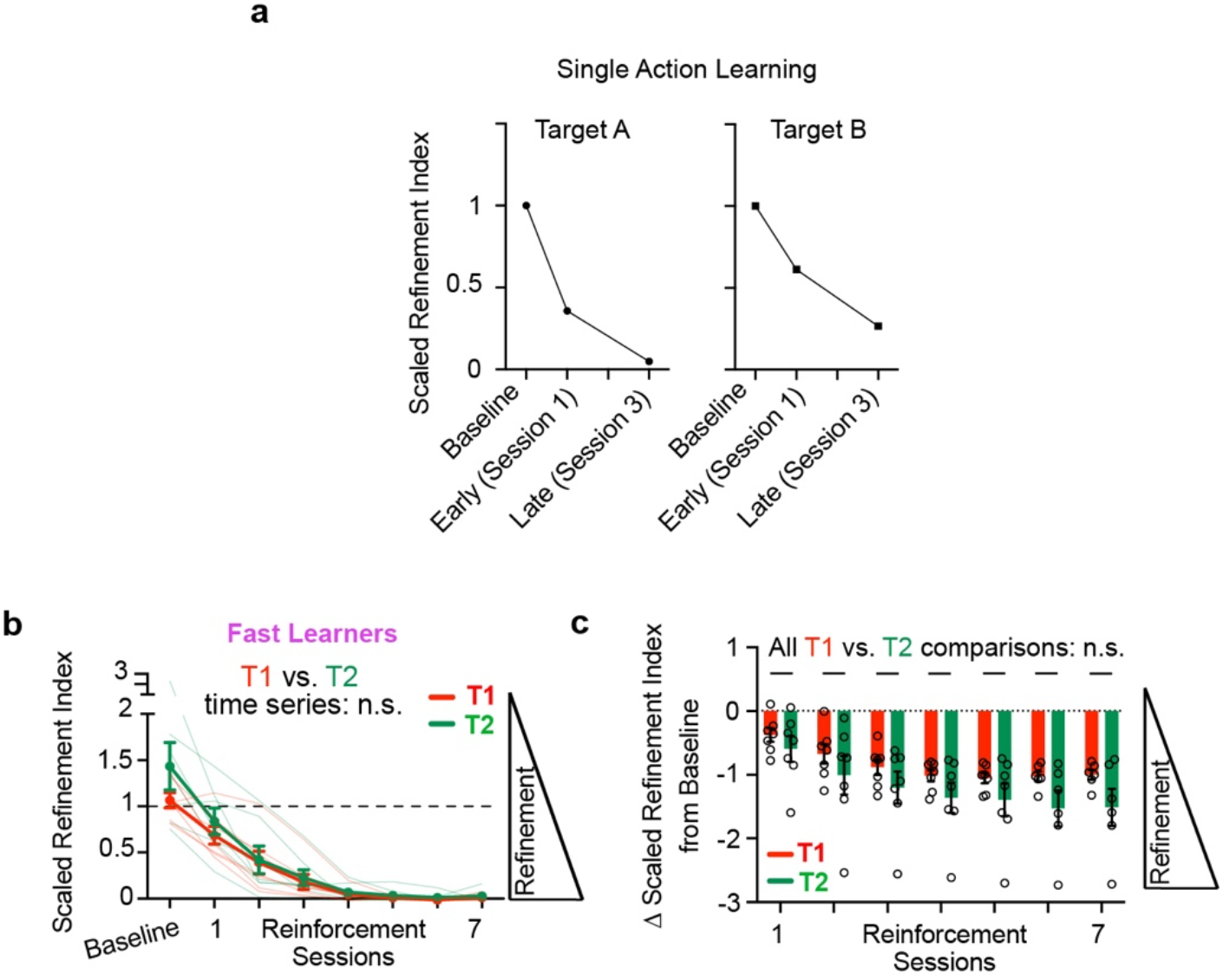
Refinement of T2 and T1 in ChR2-YFP animals. **a**, Patterns of scaled refinement index changes in Single Action Learners, extracted from normalized curves from Fig. 1k and 2e at EMD value of 0.103 (equivalent to 17th percentile - similar to the median EMD of target-related actions in sequence learning refinement analyses. **b-c,** Fast Learners (*n* = 7 mice (biological replicates)) refine T1 and T2 at similar rate. Repeated measures, mixed effects model. No significant difference between time and scaled refinement indices (raw: F(7,80) = 0.4776, p = 0.85 (**b**); Baseline-subtracted: F(6,68)=0.3351, p=0.92 (**c**)). Post-hoc Šidák test (**c**). * p < 0.05. n.s. – not significant. All summary plot elements indicate mean +/- S.E.M. See Supplementary Information for statistical/sample details.

