## Supplementary Information for "Dynamic refinement of behavioural restructure mediates dopamine-dependent credit assignment"

**Supplementary Information Guide**

Supplementary Video 1 - .mov file

Supplementary Methods – methods pertaining to the manuscript

Supplementary Notes – notes of relevance to the manuscript

Supplementary Table – statistical summary of results in the manuscript – separately attached xlsx file

**Supplementary Video 1 –** Example of an action cluster performed on 10 instances in a grey open field box, before closed loop reinforcement. Slowed down 3x.

**Supplementary Methods**

*Processing of inertial sensor data*

Clustering begins by processing accelerometer and gyroscope data to extract 4 features discriminating postural changes, movement momentum, head and head-body rotations, and total body accelerations. Feature values from 300 ms long segments of behavior were discretized into histograms, upon which pairwise similarity comparisons could be made using a Earth-Mover's Distance (EMD)[32] metric. The similarity matrix of all possible pairwise comparisons were fed into an unsupervised affinity propagation clustering algorithm[20] (Methods), identifying naturally occurring repertoire of 300 ms long behavioral clusters[21], or "actions" (Fig. 1c,Extended Data Fig. 1b). The choice to cluster actions defined as 300 ms long movements was informed by previous studies[21,41].

*Closed loop optogenetics*

Using wireless inertial sensor (Fig. 1b), we tracked behavior continuously in a white open field and used the similarity metric to match ongoing 300 ms behavioral segments to exemplars representing each mouse's repertoire of actions (Fig. 1d-e). Upon a match to a defined target action (target action A), a 25 hz, 600 ms long train of optogenetic stimulation was delivered to the VTA parabrachial pigmented area (PBP) (30-60 ms delay, Fig. 1e).

*Action dynamic classification*

Action dynamics were categorized over the course of closed loop reinforcement of a single action, action A (Fig. 1). We first sorted actions based on whether actions were significantly increased, unchanged or decreased in four stepwise, learning stage frequency comparisons that reveal whether actions were initially modulated, whether they were modulated in frequency over time, and whether they showed transient modulation in frequency (Extended Data Fig. 6).

Most actions (465 of 511 actions, 91%) were partitioned into 12 of 81 possible frequency modulation paths, with more than 10 actions per path (Extended Data Fig 6). 11 of the paths could be categorized into three meaningful types based on commonalities in action dynamics (Fig. 1m. Extended Data Fig. 6a). First, Sustain Increased dynamics indicate initially increased frequencies relative to baseline and either stable or increasing frequencies over time. Second, Transient Increase dynamics indicate initial increases in frequencies relative to baseline and subsequent decreases in frequencies. Finally, Decreased dynamics indicate initial decrease in frequencies that remain stable or continually decrease over time. Overall, 100% of target actions fell into the Sustain Increased group. Most remaining actions are of the Decreased type, while Transient Increase and Sustained Increase types comprise the second-most and least abundant dynamic types, respectively (Fig.1m).

*Multinomial Logistic Regression to predict action dynamics*

We performed multinomial logistic regression to assess whether 1- or 2-factor models best fit the observed dynamics pattern that an action would follow upon reinforcement (Fig. 3f,g). The 2-factor model fit the data best (Fig.3f) and outperformed the 1-factor models in predictions for action dynamics types (Methods; Supplementary Table). Prediction of action dynamics type with this model was significantly above chance as assessed by precision-recall curves, which is suitable for evaluating datasets with imbalanced categories[42] (Fig. 3g). The beta coefficients indicated that increased similarity to target and decreased median interval preceding target action increases the prediction of Sustained Increase and Transient Increase dynamic types relative to Decreased types (Supplementary Table).

### Detailed Statistical Procedures

#### *Single action learning*

For single action A and B learning curves repeated measures two-way ANOVA was applied (Extended Data Fig. 4b, Extended Data Fig. 9c), the data was log transformed to promote equal variance. This manipulation worsened the normality of some ChR2-YFP groups as judged by QQ-plot and normality tests above. Log-transform was chosen nevertheless because we emphasized equal variance over perfect normality of data due to the ability of ANOVA to tolerate slight deviation from normality[52]. Post hoc Tukey's multiple comparison test was used. All possible comparisons within ChR2-YFP and YFP groups (Baseline vs. Session 1/2/3 Reinforcements, Session 1 Reinforcement vs. Session 2/3 Reinforcements, Session 2 vs. Session3 Reinforcements) were considered. To detect group-specific differences between ChR2-YFP and YFP animals, the means of each time point from each group were compared between groups.

For single action A and B stimulation, repeated measures one-way ANOVA was applied (Extended Data Fig. 9d), the data was log transformed to promote equal variance. Post hoc Dunnett's multiple comparison test was used. Comparisons were made of the mean stimulations of individuals in Session 1 against those in Session 2 and 3 groups.

To test for rapid behavioral changes upon cumulative closed loop reinforcements, data were log transformed (log(Y+1). 1 was a constant to eliminate zero values). Two-way ANOVA was applied on the cumulative reinforcement vs. frequency (Fig. 1i), latency (Extended Data Fig. 5a), or average behavioral similarity (Extended Data Fig. 5b)curves to test for significant difference between ChR2-YFP and YFP animals. Post hoc Šidák test was used to perform multiple comparisons. To detect for significant changes in the tested parameters from the beginning of reinforcement, parameter values after the second cumulated reinforcement and every 5 cumulative reinforcements after the first (until 26 cumulated reinforcements) were compared to those values for the first cumulative reinforcement.

For single action A and B extinction experiments, repeated measures two-way ANOVA was applied on the maintenance, extinction (last 20 minutes) and re-acquisition frequencies of ChR2-YFP and YFP groups, respectively (Extended Data Fig. 4c; Extended Data Fig. 9g). Data was log transformed (Log(Y+1)) to improve equality of variance. Each group was treated as a family to test the hypothesis that none of the comparisons (1. maintenance period vs. extinction period 2. Extinction period vs. Re-acquisition period. 3. Maintenance period vs. Re-acquisition period) differ from each other. Post hoc Tukey's multiple comparison test evaluating all possible comparisons was used. Independence of frequencies at different time points was not assumed. For plotting, the raw untransformed data were presented (Extended Data Fig. 4c; Extended Data Fig. 9g). For comparison for difference between ChR2-YFP and YFP groups, the means of each time point from each group were compared between groups to examine group-specific differences.

For hypervolume overlap experiment (Extended Data Fig. 8), the non-parametric Wilcoxon 2-tailed test was performed to compare the percentage of transiently increased action dynamics type calculated based on all action clusters or on non-overlapping action clusters.

For single action B contingency degradation experiment, repeated measures two-way ANOVA was applied on the maintenance, contingency degradation (last 20 minutes) and re-acquisition frequencies of ChR2-YFP, laser and YFP groups, respectively (Extended Data Fig. 9i). Data was log transformed (Log(Y+1)) to improve equality of variance. Notably, the YFP group showed noticeable deviation from normality despite log transformation. However, YFP time course result tested using one-way ANOVA did not differ in conclusions from the non-parametric Friedman test. Each group was treated as a family to test the hypothesis that none of the comparisons (1. Maintenance period vs. contingency degradation period 2. Contingency degradation vs. Re-acquisition period. 3. Maintenance period vs. Re-acquisition period) differ from each other. Post hoc Tukey's multiple comparison test evaluating all possible comparisons was used. Independence of frequencies at different time points was not assumed. For plotting, the raw untransformed data were presented (Extended Data Fig. 9i). For comparison for difference between ChR2-YFP and Laser groups, the means of each time point from each group were compared between groups to examine group-specific differences in a Šidák post hoc test.

For single action A and B whole action repertoire analysis, repeated measures two-way mixed effect analysis was used (Extended Data Fig. 5c,f; Extended Data Fig. 9h,l). The two factors analyzed were percentile similarity (target, 1-100 percentile groups) and timepoint (baseline/early/mid/late). The dependent variable frequencies were log transformed (log(Y + minimum non-zero value (0.0002))) to improve equality of variance and normality of distribution. YFP control data was transformed in the same way. Post hoc Tukey's multiple comparisons test was used to compare all combinations of pairwise comparisons within percentile groups. Comparisons within each percentile similarity group were treated as a family.

For dispersion analysis, baseline and maintenance Fano factor-normalized datasets were analyzed to test whether target actions were specifically increased over the normalization condition relative to rest of actions (Extended Data Fig. 4f). To derive Fano factor values for all actions, 50 x 300 ms data points with a value of 0.02 (equivalent the detection of a single 300 ms action performance within a 15 second moving window) was appended to each analysis time window for all actions. This data transformation makes it possible to calculate Fano factor (variance/mean) for all actions. Time windows previously found to have zero action detection were found to have the lowest possible Fano factor values as would be expected. To perform bootstrap, each action in the dataset were labeled with a tag so its identity can be traced to the animal that it was from. MATLAB bootstrp function (using @mean as bootfun) was performed on the tag values of each dataset at 10,000 bootstraps. From the bootstrap result, a value from each animal was drawn at random using randi MATLAB function such that an equal number of actions sampled from each animal was collected for comparison with the original sample size of the target action dataset. Significance was determined by asking whether the original target action mean Fano factor was greater or less than the 95% confidence interval of the bootstrap distribution.

The nonparametric Kruskal-Wallis test was performed to compare average number deviations from initial modulation status of initially increased, unchanged, and decreased groups (Extended Data Fig. 7d). Dunn’s post hoc test was used to make multiple comparisons of all permutations.

Repeated measures, two-way ANOVA was performed to test the hypothesis that there is no difference in baseline-subtracted moving mean frequencies between action A and action B during early and late time bins following the start of reinforcement for either action (Fig. 2i-j). Data was normally distributed. Post hoc Šidák’s multiple comparison test was conducted comparing action A and B values from each specified time bin.

Mixed effects model was performed to test the hypothesis there is no difference in temporal profile of inter-target action interval changes between ChR2-YFP and YFP animals for action A and B reinforcements (Fig. 3b). Data was log transformed to improve normality and equal variance. A single outlier was removed from the YFP, Reinforce B dataset, because the median inter-action interval was calculated from a single value from Session 6 (Day 2 of Reinforce B). Post hoc Tukey’s multiple comparison test evaluating all possible comparisons was used. Only specific comparisons of interest were show in the plots. For plotting, the raw untransformed data was presented (Fig. 3b).

Permutation test was used to test the hypothesis that there is no difference in the means of Time to Target or Action Similarity values in pairwise comparisons between action dynamic types (Fig. 3d-e). 10,000 permutations were tested. Bonferroni p adjustment was performed for each group of comparisons (3 comparisons per group). Statistical significances of lower than alpha = 0.05 are maintained even when considering paired comparisons from both Similarity and Time to Target groups as a multiple comparison experiment.

Two-way ANOVA was used to test the hypothesis that deviances between Real and Shuffled regression conditions were significantly different (Fig. 3f). Post-hoc Dunnett test was used to perform multiple comparisons between mean deviances of double factor regression model vs. single-factor models.

Permutation test was used to test the hypothesis that there is no difference in the means of true positive action dynamic type predictions between regression models created using median time preceding target or action similarity with the model created using both factors (Fig.3f(III) tab in Supplementary Table). 10,000 permutations were tested. Bonferroni p adjustment was performed (6 comparisons).

Permutation test was used to test the hypothesis that there is no difference in the means of area under the precision-recall curve (PRC) curve between regression models created using Real data or Shuffled data or the mean of the proportion of positives for each action dynamic category tested (Baseline) (Fig. 3g). 10,000 permutations were tested. Bonferroni p adjustment was performed for each group of comparisons (2 comparisons per group). Statistical significances of lower than alpha = 0.05 are maintained even when considering paired comparisons from all three groups as a multiple comparison experiment.

Mixed-effect model was performed to test the hypotheses that there is no difference in temporal profile of baseline normalized frequencies of action transitions between Forward and Retrospective conditions in ChR2-YFP and YFP animals for action A reinforcement (Fig. 3j). Forward and Retrospective conditions comprise of sliding 1.2 second windows occurring after and before an initial start window, respectively. The initial start window is chosen such that the start of stimulation upon target reinforcement occurrence is coincident with the second of three action transitions in this particular sliding 1.2 second window. Data was log transformed (Log(Y + 0.001)) to improve normality and equal variance. Sphericity was not assumed, and Geiser-Greenhouse correction was applied. Post-hoc Šidák multiple comparisons test evaluating matching Forward and Retrospective data points was used. For plotting, the raw untransformed data was presented (Fig. 3j).

*Two Action Section*

To test that the mean frequency (Fig. 4e), T1→T2 median interval (Fig. 4i), and refinement indices (Fig. 4k) were significantly changed in ChR2-YFP Criterion and YFP animals over time, repeated-measures one-way ANOVA was used. Mean frequency and refinement index data were log-transformed to promote normality. T1→T2 median interval data showed normality without transformation. Post-hoc Šidák test was used to perform multiple comparisons between means of different measurements of baseline and reinforcement portions of session 1 as well as between reinforcement portion of session 1 and last session for each animal. As an exception, the last session for one of the YFP animals could not be used for median T1→T2 interval because no triggers were induced. The second-to-last session was thus used for this YFP animal.

To test for statistical significance as to whether reinforced action pairs differ in initial inter-trigger intervals (Extended Data Fig. 11f), we examined the first 11 bins (totaling to 2400 cumulated reinforcements) because these were reached by all individuals by the end of learning. A Y = log(Y + 1) transformation was applied to improve normality of data. 2-way ANOVA was performed, with reinforcement bins representing the Time factor. Slow/Fast learners are divided into two groups. Post-hoc Sidak multiple comparisons analysis was then performed comparing the scaled frequencies of Slow and Fast learners at matching reinforcement bins.

To test for significant differences in the independent variables in Extended Data Figure 11b-d, we conducted 2-tailed, unpaired t-test assuming equal variance.

We controlled for non-specific increase in sequences by comparing whether B→A sequence frequency progression over single action reinforcement for target action A were significantly different from T1→T2 frequency progression over two action sequence reinforcement for T1→T2, two-way ANOVA was applied (Extended Data Fig. 12n). The data was transformed with a log(data+1) formula. The progression of learning from baseline to reinforcement session 1 and last session formed the Time factors. Single action and action sequence reinforcement samples formed the Group factors. Post hoc Dunnett's multiple comparison test was used to test whether frequencies change from baseline conditions for each group. Comparisons were made on the mean frequencies of B→A or T1→T2 sequences between baseline and session1 or last session.

To test that the mean frequency was significantly changed in ChR2-YFP Criterion animals over the course of extinction, repeated-measures one-way ANOVA was used (Fig. 4g). Frequencies (triggered per minute) from the last 5 minutes of maintenance, extinction and re-acquisition conditions were analyzed. Data were log-transformed to promote normality. Post-hoc Šidák test was used to perform multiple comparisons between means of all possible pairwise comparison between protocol conditions.

For Fast vs. Slow Learner median time interval comparisons at different learning points, repeated measures two-way ANOVA was applied on values from open field, session of Turning Points, first session above criterion frequency, and last session of Fast and Slow Learner groups, respectively (Fig. 5d, Extended Data Fig. 13c). Data was normalized by log transformation. Each group was treated as a family to test the hypothesis that none of the comparisons differ from each other (Supplementary Table). Post hoc Tukey's multiple comparison test evaluating all possible comparisons was used. For plotting, the raw untransformed data were presented (Fig. 5d, Extended Data Fig. 13c). For comparison for difference between Fast and Slow Learner groups, the means of each time point from each group were compared between groups to examine group-specific differences.

For preferential refinement of T1 vs. T2 analyses. Scaled refinement values of T1 and T2 actions for each animal were log-transformed (Log(Y+1)) to promote normality of data. Repeated measures, mixed effects analyses were conducted on Slow Learners (Fig. 5f-g). First, the null hypothesis that the time course of scaled refinement indices did not differ between T1 and T2 actions was tested (Supplementary Table), followed by post-hoc Šidák test to examine whether scaled refinement indices changed from Starting Point to subsequent conditions of Turning Point, Session of Criterion Frequency, or Last Session (Fig. 5f). Second, the null-hypothesis that the time course of Starting Point-subtracted scaled refinement indices did not between T1 and T2 actions were tested (Supplementary Table), followed by post-hoc Šidák test to examine whether normalized scaled refinement indices of T1 and T2 differ between each other in each examined learning stage (Fig. 5g). For Fast Learners, similar procedures were performed (Extended Data Figure 14b-c).

For odds ratio statistics, two-tailed, paired Wilcoxon test was used to test the hypothesis that there is no significant difference between the odds ratios at open field→Turning Point and Turning Point→session of criterion frequency periods (Fig. 5e).

For T1 probability rank change across time bins from T2 trigger, repeated measures, two-way ANOVA was used to test the hypothesis that the time course progression of T1 parameters across learning stages were significantly different between first occurrences of T1 preceding and following T2 trigger (Supplementary Table). Post hoc Šidák multiple comparisons test was performed to test the null hypothesis that the total T1 parameter is not different between Starting Point values and Turning Point, session of criterion frequency, or last session (Fig. 5i).

**Supplementary Notes**

*Hypervolume analysis in single action learning*

As t-SNE do not include higher dimensional information, we analyzed overlap of action cluster hypervolumes in higher dimensional space (Methods). The range of maximum proportion of an action cluster hypervolume overlapping with target action hypervolume per animal is very low, on the order of 1*10^-5 to 1*10^-30 (Value of 0 is no overlap. Value of 1 indicate perfect overlap between hypervolumes). The percentage of transiently increased actions per animal remain unchanged even after accounting for overlapping action clusters (Wilcoxon 2-tailed test: $p = 0.25$, n= 15 ChR2-animals) (Extended Data Fig. 8c). Thus, transiently increased dynamics remain the predominant fate of initially reinforced actions.

*Different responses of animals upon naïve reinforcement and target action changes*

Whereas naïve animals responded to initial reinforcements for target action A by significantly increasing action A performance relative to the non-target action B (Fig. 2g,i,left graph), animals with a history of reinforcement on action A animals responded to initial reinforcements of action B by increasing non-target action A performance (Fig. 2g,i,right graph). This trend reverses later such that target action B becomes significantly increased over the non-target action A (Fig. 2g,i,right graph). YFP control animals showed no such trends (Fig. 2h,j). Thus, DA reinforcement does not simply reinforce the recently performed, temporally contiguous action, but trigger previously credited actions in the face of a new action-reward contingency that is not yet learned. This suggest again that animals learned the contingency between action performance and DA release.
